# Placental inflammation leads to abnormal embryonic heart development

**DOI:** 10.1101/2022.03.10.482611

**Authors:** Eleanor J Ward, Serena Bert, Silvia Fanti, Neil P Dufton, Kerri M Malone, Robert T Maughan, Fabrice Prin, Lia Karina Volpato, Anna Paula Piovezan, Mauro Perretti, Federica M Marelli-Berg, Suchita Nadkarni

## Abstract

Placental and embryonic heart development occurs in parallel, and these organs have been proposed to exert reciprocal regulation during gestation. Poor placentation has been associated with congenital heart disease (CHD), an important cause of infant mortality. However, the mechanisms by which altered placental development can lead to CHD remain unresolved. In the current study we show that neutrophil-driven placental inflammation leads to inadequate placental development and loss of barrier function. Consequently, placental inflammatory monocytes of maternal origin become capable to migrate to the embryonic heart and alter the normal composition of resident cardiac macrophages and cardiac tissue structure. This cardiac impairment continues into postnatal life, hindering normal tissue architecture and function. Finally, we demonstrate that tempering placental inflammation can rescue this fetal cardiac defect and is sufficient to promote normal cardiac function in postnatal life. Taken together, our observations provide a mechanistic paradigm whereby neutrophil-driven inflammation in pregnancy can preclude normal embryonic heart development as a direct consequence of poor placental development.

## Introduction

The placenta is a specialized organ that acts as a tight barrier to regulate the transfer of oxygen and nutrients to the developing fetus, whilst preventing passage of harmful pathogens and cells. Placental and embryonic heart development occur in parallel, suggesting that the two organs influence each other’s development^1^. The heart is one of the first organs that develops in the embryo, with sequences in cardiac development conserved in both mouse and human^2^. Clinical studies suggest a strong association between placental dysfunction and congenital heart disease (CHD)^3, 4^, with poor trophoblast invasion, and aberrant oxygen and nutrient transfer leading to poor cardiac development^5, 6^. CHD is an important cause of infant mortality associated with approximately 35% of infant deaths^7^. Despite causal factors being hypothesized, the pathogenesis of CHDs has not been resolved to-date.

The role of tissue-resident macrophages in promoting normal organogenesis is well established ^8,9^. At around E8.5, yolk sac-derived erythro-myeloid progenitors (EMP) migrate to the developing embryo in a chemokine-dependent manner. These EMPs develop into premacrophages expressing CX_3_CR1, Kit and CSFR1 and seed various organs including brain, liver, and heart, where they persist into adulthood through a process of self-renewal ^10, 11^. In the heart, CX_3_CR1^+^CCR2^−^ macrophages migrated from the yolk sac promote angiogenesis and regulate coronary vascular development^12^, and exert reparative functions in adult cardiac tissue ^13^. This contrasts with blood monocyte-derived CCR2^+^ macrophages that promote inflammation in the adult heart ^13, 14, 15^.

In the current study, we report that circulating pro-inflammatory neutrophils during gestation can cause placental inflammation and the breakdown of placental barrier function. This pathological neutrophil-driven placental inflammation allows migration of inflammatory maternal monocytes to the embryonic heart, which, in turn, lead to abnormal fetal cardiac development with inadequate cardiac function in postnatal life.

## Results

### A Neutrophil-driven model of placental inflammation

We have previously shown that depletion of maternal neutrophils at E4.5 and E7.5 results in abnormal placental development, with shallow trophoblast invasion and poor spiral artery remodeling at E12.5 ^16^. Following antibody depletion, neutrophils return to the circulation within 72hrs ^17^ and these cells now present an activated, pro-inflammatory phenotype, characterized by high CXCR2 and CD114 (G-CSFR) expression. No differences are observed in circulating proinflammatory CCR2^+^ monocytes (Figure 1 A and B). We next investigated the placental environment in more detail. Maternal neutrophil depletion resulted in smaller placenta weights and shallow trophoblast invasion (Figure S1B and S1C), coupled with an exaggerated TNΦ-α concentration within the placental tissue, but not the circulation (Figure 1C). There was no overall difference in total CD45^+^ leukocyte numbers (Figure 1D and Figure S1D), nor in numbers of CD3^+^ T cells and NK cells (Figure S1E). Although placentae from neutrophil-depleted mothers displayed no overall difference in the number of neutrophils compared to their control counterparts (Figure 1E), their placental neutrophils displayed an activated phenotype with high expression of TNF-α, CXCR2, CD114 (G-CSFR), and MMP9 (Figures 1F and G). Therefore, neutrophil depletion in pregnant mice induces *neutrophil-driven placental inflammation* (referred to hereafter as NDPI).

**Figure 1:**
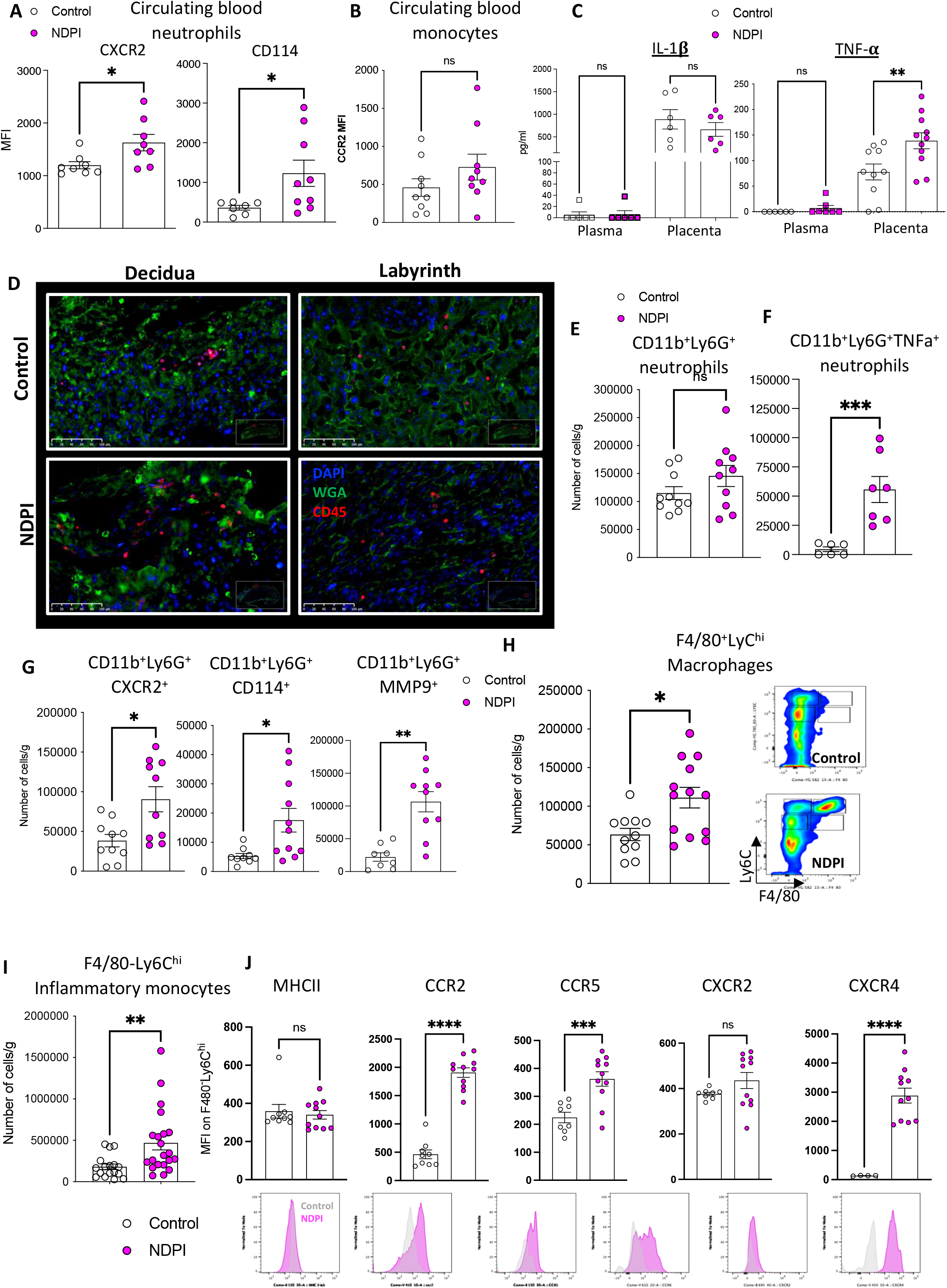
A Neutrophil-driven placental inflammation model. Neutrophils were depleted at day 4.5 and 7.5 of pregnancy using αLy6G. Mice were sacrificed at E14.5 of pregnancy and placentae harvested to assess the structure and immune composition. Isotype control treated (white) and neutrophil depleted (NDPI) (pink). (A) Expression of activation markers by maternal blood neutrophils (B) CCR2 expression on maternal blood monocytes (C) ELISA showing the concentration of IL-1β and TNF-α in plasma and placenta digest supernatants. (D) Immunofluorescent staining of placentae for CD45 (red) WGA (green) and cell nuclei with DAPI (blue). Expression is shown in the decidual layer (left) and labyrinth layer (right). (E-G) Neutrophil subpopulations from placentae analyzed by flow cytometry expressed as absolute cell number per gram of tissue. (E) CD11b^+^Ly6G^+^ neutrophils, (F) CD11b^+^Ly6G^+^ TNFα^+^ neutrophils, (G) neutrophils expressing CD11b^+^Ly6G^+^ CXCR2, CD11b^+^Ly6G^+^ CD114+ and CD11b^+^Ly6G^+^ MMP9^+^. (H) Macrophages from placentae analyzed by flow cytometry expressed as absolute cell number per gram of tissue. (I) Inflammatory monocytes from placentae analyzed by flow cytometry expressed as absolute cell number per gram of tissue. (J) F4/80^−^Ly6C^hi^ populations from placentae were analyzed for the expression of MHCII, CCR2, CXCR2, CCR5 and CCR6, expressed as median fluorescent intensity. Representative histograms are shown below the quantification (control= grey, NDPI =pink). Each symbol represents an individual mouse from different pregnancies and statistical significance was tested by unpaired Student’s t-test. ns = not significant, *p≤0.05, ** p≤0.01, ***p≤0.001 **** p≤0.0001. In all cases, data are mean ± SEM. MFI, median fluorescence intensity

NDPI was also defined by the presence of activated F4/80^+^Ly6C^hi^ macrophages (Figure 1H), with no difference in the number of these macrophages making TNF-α between both groups (Figure S1F), suggesting that neutrophils are the likely source of increased TNF-α in NDPI. We also observed that intra-placental monocytes displayed an inflammatory phenotype compared to control placentas (Figure 1I). These F4/80^−^Ly6C^hi^CCR2^+^ monocytes also expressed higher levels of CCR2, CCR5 and CXCR4 on their surface, but not MHC II or CXCR2 (Figure 1J) compared to monocytes from control placentas. Lack of difference in total leukocyte numbers between placentas from control and NDPI pregnancies (Figure S1D), coupled with no overall change in the inflammatory status of circulating maternal monocytes, indicated that the phenotype of placental macrophages and monocytes is not due to an influx from the maternal circulation, but rather to *in-situ* activation. Thus, we hypothesized that TNF-α-producing neutrophils within the placental tissues regulate the phenotype of placental monocytes and macrophages. To challenge this hypothesis, we isolated placental neutrophils from control and NDPI pregnancies and co-cultured them with naïve splenic monocytes from non-pregnant aged-matched mice. These co-cultures showed the induction of higher proportions of inflammatory F4/80^+^Ly6C^hi^ macrophages following monocyte exposure to NDPI neutrophils, mirroring the inflammatory phenotype observed in NDPI *in vivo* (Figure S1G and S1H).

### Neutrophil-driven Inflammation promotes a breakdown in placental tissue barrier

To further investigate the features of NDPI we undertook bulk RNA sequencing of E14.5 placental tissues from control and NDPI pregnancies. In NDPI we observed downregulation of 329 genes, out of 357 genes in total to be significantly changed (false detection rate, FDR, corrected P<0.05) (Figure 2A). Pathway analyses identified post-translational protein phosphorylation, collagen trimerization and nutrient transport as the top pathways (Figure 2B). Downregulation of the latter pathway included lipid, glucose and carbohydrate metabolism, as well as genes involved in amino acid transport (Figure S2A) and suggests NDPI may impact on the transfer of nutrients to the fetus, with likely fetal growth restriction as we previously reported^16^.

**Figure 2:**
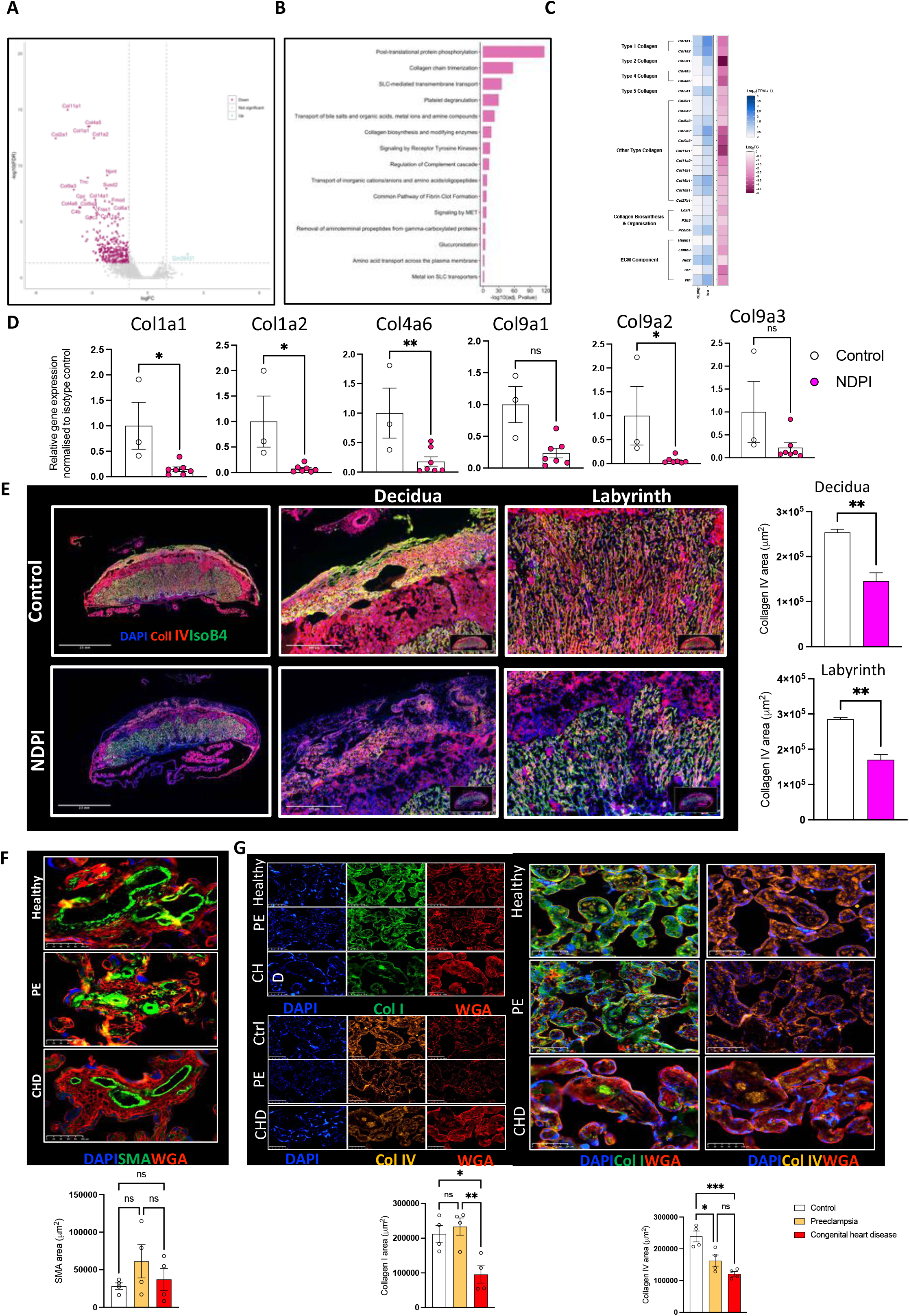
Neutrophil-driven Inflammation promotes a breakdown in the placental tissue barrier. (A-E) Neutrophils were depleted at day 4.5 and 7.5 of pregnancy using αLy6G. Mice were sacrificed at E14.5 of pregnancy and placentae harvested. Isotype control treated (white) and neutrophil depleted (NDPI) (pink). (A-C) RNA sequencing from placentae. (A) Volcano plot showing differentially expressed genes in control versus NDPI placentae (B) Top 15 modulated pathways in placentae of NDPI mice when compared to isotype control (C) Heat map showing gene expression of extracellular matrix components of NDPI or control placentae (D) RT-PCR showing the gene expression of Col1a1, col1a2, col4a6, col5a1, col9a1, col9a2 and col9a3 (E) Immunofluorescent images of control or NDPI murine placentae, stained with DAPI (blue) collagen IV (red) isolectin B4 (red). Magnified images of the decidua and labyrinth of the placenta and graphs showing the quantification of Col IV staining. (F-G) Immunofluorescent staining of term placentae from women with healthy, preeclamptic pregnancies or pregnancies carrying babies with congenital heart disease for (F) SMA (green) (G) Col I (green) (top panel) Col IV (orange) bottom panel. In all panels WGA staining in red and cell nuclei are stained with Dapi (blue). Quantification of area stained with SMA, Col I and Col IV shown in panel below images Each symbol represents an individual sample from different pregnancies and statistical significance was tested by unpaired t-test. ns = not significant, *p≤0.05, ** p≤0.01, **p≤0.001. In all cases, data are mean ± SEM.

We focused our attention on collagen genes, since these extracellular matrix (ECM) components are important for placental tissue integrity^18^. Further analyses of the collagen genes revealed that of the 44 collagen genes within the mouse genome, 17 were significantly downregulated in NDPI (Figure S2B): including *Col1a1 and Col1a2, Col2a1, Col4a6, Col9a2* and *Col9a3, Col11a1, Col11a2* and *Col11a3* (Figure 2C and 2D). Other significantly downregulated ECM genes included those required for collagen biosynthesis, prolyl3-Hydrolase 3 (*P3H3)*, and procollagen C-endopeptidase enhancer 1 (*Pcolce*)^19, 20^.

Immunofluorescent staining for two collagens (type I and IV) important for placental tissue integrity^18^ demonstrated lower expression of collagen I in the decidua of inflamed placentae, but no difference in the labyrinth, when compared to control placentae (Figure S3A). For collagen IV, significant reductions were quantified in both decidua and labyrinth of inflamed placentae (Figure 2E). Of note, collagen IV is required not only for placental tissue integrity for but also for the invasive properties of trophoblasts^21^, and thus the reduced collagen IV expression we report here, may explain the shallow trophoblast invasion displayed in NDPI settings.

We next sought to test whether this collagen phenotype could be observed in human tissues in two types of pregnancy complications that affect placental development (Table 1). Firstly, we assessed placental tissue from preeclamptic pregnancies (PE), chosen because there is evidence for an activated neutrophil phenotype in these patients^16, 22^. Second, we analyzed placental tissue from pregnancies associated with fetuses that have CHD in the absence of maternal PE. We observed a significant reduction in collagen IV expression in trophoblasts from both PE *and* CHD placentae compared to healthy placentae (Figure 2G, red). This suggests a compromised trophoblast invasion in these placentae since collagen IV is important for normal trophoblast invasion^21^. There was a 2-fold attenuation in trophoblast collagen I expression in placentae of CHD fetuses when compared to control and PE placentae (Figure 2G, green). Together, these data indicate NDPI has a negative functional impact on the placental support structure.

**Table 1:** Demographics of pregnant patients from which placental tissue was assessed. Patient demographics of placentae characterized in Figure 2F. Words in Bold highlight each patient demographic. CHD= fetal congenital heart disease

| **Pregnancy type** | **Age of Mother** | **Parity** | **Gestation (weeks)** | **Blood pressure** | **Mode of delivery** |
| --- | --- | --- | --- | --- | --- |
| Healthy | 18 | 0 | 41 | Within normal range | Vaginal |
| Healthy | 28 | 0 | 41 | Within normal range | Vaginal |
| Healthy | 31 | 0 | 39 | Within normal range | Vaginal |
| Healthy | 32 | 3 | 40 | Within normal range | Vaginal |
| Healthy | 40 | 5 | 37 | Within normal range | Vaginal |
| Pre-eclampsia | 30 | 2 | 37 | 190/110 | C-section |
| Pre-eclampsia | 22 | 0 | 37 | 171/106 | C-section |
| Pre-eclampsia | 32 | 2 | 36 | 178/113 | C-section |
| Pre-eclampsia | 32 | 2 | 38 | 190/120 | C-section |
| Pre-eclampsia | 22 | 0 | 40 | 192/101 | Vaginal |
| Fetal CHD:<br>D-transportation of great arteries | 39 | 0 | 39 | Within normal range | C-section |
| Fetal CHD:<br>Ventricular septal defect | 28 | 0 | 38 | Within normal range | C-section |
| Fetal CHD:<br>Tetralogy of Fallot | 33 | 0 | 35 | Within normal range | C-section |
| Fetal CHD:<br>Ventricular septal defect | 36 | 0 | 40 | Within normal range | C-section |

### Placental inflammation leads to poor embryonic cardiac development

Because of the discussed association between physiological placental status and embryo heart, we investigated whether NDPI impacts on embryonic heart development. Initial observations revealed an abnormal heart development in the gross structure of embryonic hearts at E14.5 from NDPI pregnancies, with significantly thinner left ventricular walls compared to control (Figures 3A,3B). Between E9.0 and E9.5, primitive cardiomyocytes within the ventricular wall form finger-like projections called trabeculae that are lined by endocardial cells. As cardiac development progresses, the ventricles undergo a switch from a mostly trabecular to a compacted state where cardiomyocytes compact and increase ventricular wall thickness; this process is essential for normal heart function. Dysregulation of this switch can cause hypertrabeculation, leading to a congenital cardiomyopathy named left ventricular non-compaction or LVNC ^23, 24^. 3-D imaging using high resolution episcopic microscopy revealed increased trabeculation of the ventricular walls of hearts from NDPI embryos, compared to controls (Figure 3C). Recent studies have demonstrated that dysregulation of coronary endothelial cells promotes the development of LVNC ^23, 25, 26^. With this in mind, we interrogated the status of the endothelial cells (ECs) within the embryonic heart. Flow cytometric and immunofluorescent staining revealed reduced numbers of CD31^+^ EC in NDPI embryonic hearts compared to controls (Figure 3D), which was due to attenuated proliferation as assessed by *in vivo* BrdU incorporation. Moreover, we observed reduced expression of key regulators of angiogenesis including ICAM-1, VCAM-1, endoglin, and thrombospondin on these cells (Figure 3E), indicating that NDPI impedes normal embryonic heart development by potentially hindering embryonic heart vascularization.

**Figure 3:**
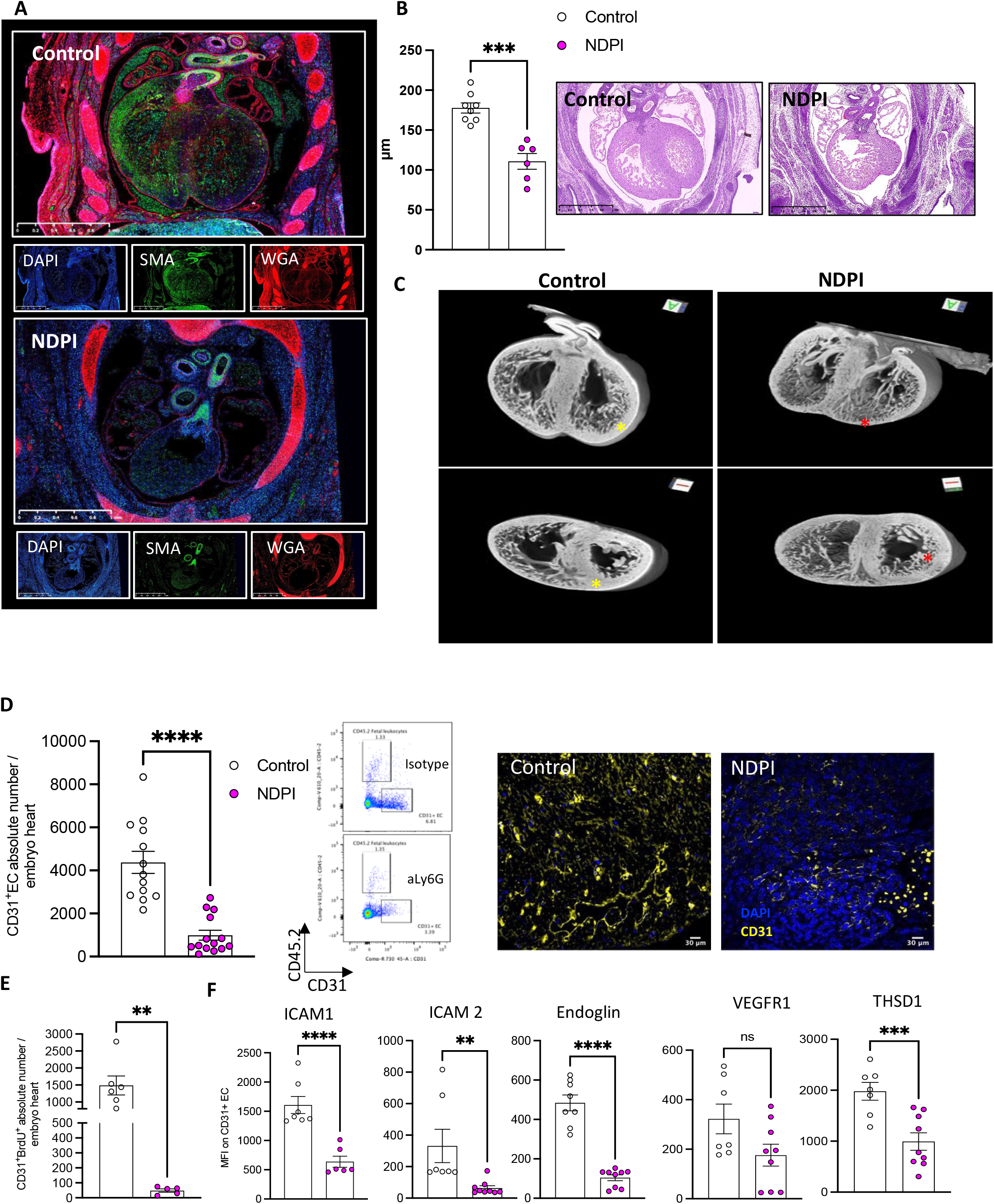
Placental inflammation leads to poor embryonic cardiac development. Neutrophils are depleted at day 4.5 and 7.5 of pregnancy using αLy6G. Mice are sacrificed at E14.5 of pregnancy and embryos harvested. Isotype control treated (white) and neutrophil depleted (NDPI) (pink). (A) Immunofluorescent images of E14.5 embryo heart of cross section. Sections were stained with DAPI (blue) WGA (red) SMA (green). (B) H&E stained cross sections of embryo heart and quantification of left ventricle wall thickness (C) High-resolution episcopic microscopy images of asterix denotes trabeculations in the ventricle wall. (D) Quantification of CD31+ endothelial cells in embryo hearts and representative FACs plots, showing CD31+ cells and CD45.2+ cells of fetal origin. Immunofluorescent staining of embryo hearts for CD31 (yellow) and cell nuclei with DAPI (blue). Scale bar 30μM. (E) Quantification of the *in vivo* uptake of BRDU into endothelial cells in embryo hearts. (F) CD31+ endothelial cells from hearts of E14.5 embryos were analyzed for the expression of ICAM1, ICAM2, Endoglin, VEGFR1 and THSD1. Quantification showing the median fluorescent intensity (MFI) of the corresponding surface marker. Each symbol represents an individual mouse and statistical significance was tested by unpaired Student’s t-test. ns = not significant, *p≤0.05, ** p≤0.01, ***p≤0.001 **** p≤0.0001. In all cases, data are mean ± SEM.

### Embryonic hearts from NDPI pregnancies present increased numbers of maternal leukocytes with a proinflammatory phenotype

The data so far indicate NDPI leads to an upregulation in inflammatory monocyte phenotype within the placental tissue; this is coupled with a dysregulation in the tight placental tissue barrier and abnormal embryonic heart development associated with reduced angiogenesis. To test whether immune cells from inflamed placentas have the capacity to directly shape embryonic heart development, we utilized the CD45.1/CD45.2 system, which allowed discrimination of immune cells of maternal or fetal origin (see Methods). Maternal cells were identified as CD45.1^+^CD45.2^−^ (referred to hereafter as maternal), whereas fetal cells were identified as CD45.1^+^CD45.2^+^ (referred to hereafter as fetal). Flow cytometric analyses revealed a near 4-fold increase in the absolute number of maternal cells in the embryonic heart of NDPI, compared to control, with no significant differences observed in maternal cells in the fetal liver (Figure 4A). Phenotypic analyses of maternal leukocyte subsets revealed presence of T cells, NK cells, neutrophils, monocytes with the predominant maternal cell type in the embryo heart being Ly6C^hi^F4/80^+^ macrophages (Figure 4B). Detailed phenotypic analyses of maternal F4/80^+^Ly6C^hi^ macrophages showed no difference in MHC II expression between control and NDPI pregnancies, yet we quantified significant differences in expression of proinflammatory chemokine receptors CCR2, CCR5 and CXCR4 (Figure 4C). This chemokine receptor expression profile was akin to that of placental inflammatory F4/80^−^Ly6C^hi^ monocytes from NDPI pregnancies, adding support that these maternal cells in the embryo heart originated from the placenta (Figure 1J).

**Figure 4:**
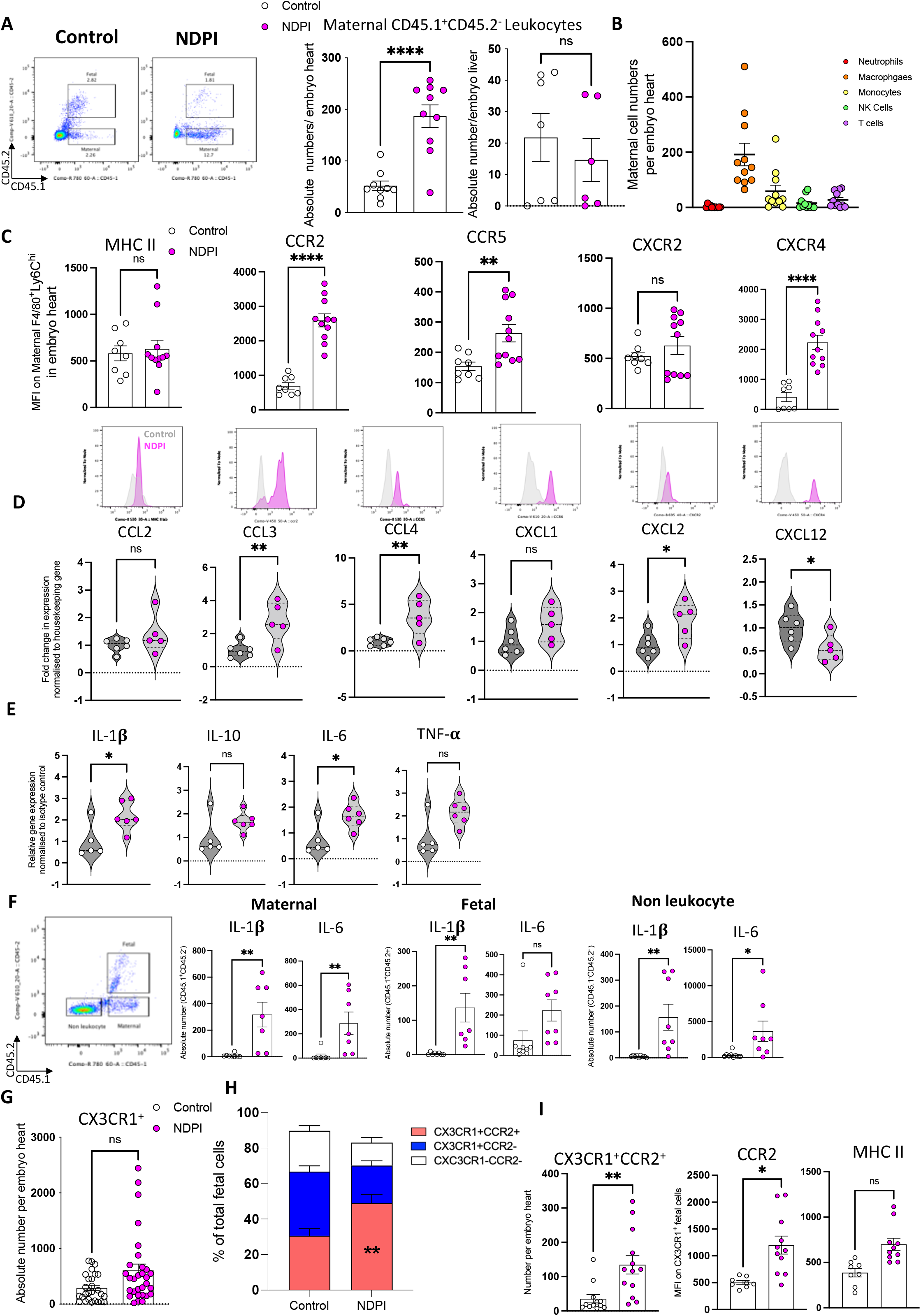
Embryonic hearts from NDPI pregnancies have an increased number of maternal leukocytes and a proinflammatory phenotype. Neutrophils are depleted at day 4.5 and 7.5 of pregnancy using αLy6G. Mice are sacrificed at E14.5 of pregnancy and hearts dissected from harvested embryos. Control (white) and neutrophil depleted (NDPI) (pink). (A) Flow cytometry plots of leukocytes in E14.5 fetal hearts from isotype and αLy6G treated pregnancies; CD45.1^+^ CD45.2^−^ cells are of maternal origin and CD45.1^+^CD45.2^+^ cells are of fetal origin. Graphs show quantification of number of maternal cells per embryo heart or embryo liver. (B) Flow cytometry quantification of different maternal leukocyte subsets in embryo hearts. (C) Median fluorescent intensity of MHCII, CCR2, CXCR2, CCR5 and CCR6 on maternal F4/80+Ly6Chi cells found in embryo hearts. Lower panel, representative histograms. (D-E) Violin plot showing gene expression of chemokines (D) and cytokines (E) in embryo hearts from control and NDPI pregnancies normalized to control (F) Representative FACs plot showing non leukocytes, maternal leukocytes (CD45.1^+^ CD45.2^−^) and fetal cells (CD45.1^+^CD45.2^+^) in embryo heart. Graphs show quantification of number of cells in maternal or fetal origin leukocytes or from non-leukocytes expressing Il-1β and IL-6. (G) Quantification of total number of cells found in embryo hearts. (H) Proportion of CX3CR1^+^CCR2^+^, CX3CR1^+^CCR2^−^ and CX3CR1^−^CCR2^−^ cells per embryo heart. (I) Quantification of fetal heart inflammatory CX3CR1^+^CCR2^+^ cells. Median fluorescent intensity of MHCII, CCR2 expression on fetal macrophages. Each symbol represents an individual mouse and statistical significance was tested by unpaired Student’s t-test. ns = not significant, *p≤0.05, ** p≤0.01, ***p≤0.001 **** p≤0.0001. In all cases, data are mean ± SEM. MFI, median fluorescence intensity

q-PCR analyses revealed significant increases in gene expression for both CCL3 and CCL4, ligands for CCR5, as well as CXCL2, with a reciprocal attenuation in CXCL12 (Figure 4D) in the embryonic heart from NDPI pregnancies, compared to control; these changes were organ-specific, since no differences were observed in embryonic livers (Fig S4A) and suggest targeted migration of maternal cells specifically to the embryonic heart. These analyses also revealed a significant augmentation in both IL-6 and IL-1□ in NDPI embryonic hearts, but not TNF-α or IL-10 (Figure 4F). Intracellular flow cytometry identified a significant up-regulation in the absolute number of maternal, fetal, and non-leukocytes expressing IL-1□ in NDPI hearts, and only maternal macrophages and non-leukocytes expressing higher levels of IL-6 (Figure 4G). Altogether, these data indicate that a mother-driven proinflammatory environment within the embryonic heart interferes with normal development of the organ.

To mechanistically challenge this hypothesis, we focused on resident fetal cardiac macrophages. In the fetal heart, CX_3_CR1^+^ are resident macrophages that originate from the yolk sac. These comprise two populations: first a CCR2^−^ macrophage population that exert pro-angiogenic functions during cardiac development ^14, 15, 27^ and second a CCR2^+^ macrophage population that have a short life-span within the embryonic and neonatal heart, with currently unknown function ^13, 27^. Closer examination of CX_3_CR1^+^ fetal macrophages did not show differences in absolute numbers between control and NDPI pregnancies (Figure 4H). However, an increase in percentage of inflammatory CX3CR1^+^CCR2^+^ phenotype was quantified in NDPI embryonic hearts compared to their control counterparts was observed (Figure 4I), with a ~4-fold increase of CX3CR1^+^CCR2^+^ fetal macrophages per heart. These cells doubled their CCR2 surface expression with no difference for MHC II expression (Figure 4J). The relative reduction of CX3CR1^+^CCR2^−^ heart-resident fetal macrophages may underlie the reduced endothelial cell proliferation in NDPI embryonic hearts.

To gain a deeper understanding of leukocyte composition and phenotype in the embryonic hearts, we undertook single cell RNA-sequencing (scRNA-seq). CD45^+^ leukocytes were isolated from E14.5 embryonic hearts from control and NDPI pregnancies. UMAP analyses revealed heterogenous leukocyte populations with a mixture of both myeloid and lymphoid cells present (Figure 5A, 5B). Overlay of UMAPs from control and NDPI identified differences in the cluster that we identified as resident fetal macrophages, expressing genes associated with the cells, including *Cx3cr1, Ccr2, Folr2, C1qc* (Figure 5C). Further analyses of embryonic leukocytes within the hearts revealed less resident fetal macrophages (−27% of total leukocyte population) associated with an increase in leukocytes of maternal origin (+14.4% of the total leukocyte population), when comparing control and NDPI groups (Figure 5D). Through EnrichR analysis ^28^, we identified the fetal resident macrophage cluster. In control embryonic hearts, this cluster was selectively enriched of transcripts of myeloid origin, while the same cluster from NDPI embryo hearts displayed a mixture of myeloid *and* lymphoid gene transcripts (Figure 5E). Whether this is due to seeding from cells of maternal origin or because heart proximity to the thymus requires further investigations, though it suggests that dysregulation of the maternal innate immune response during early pregnancy impacts on the innate immune response during embryonic development.

**Figure 5:**
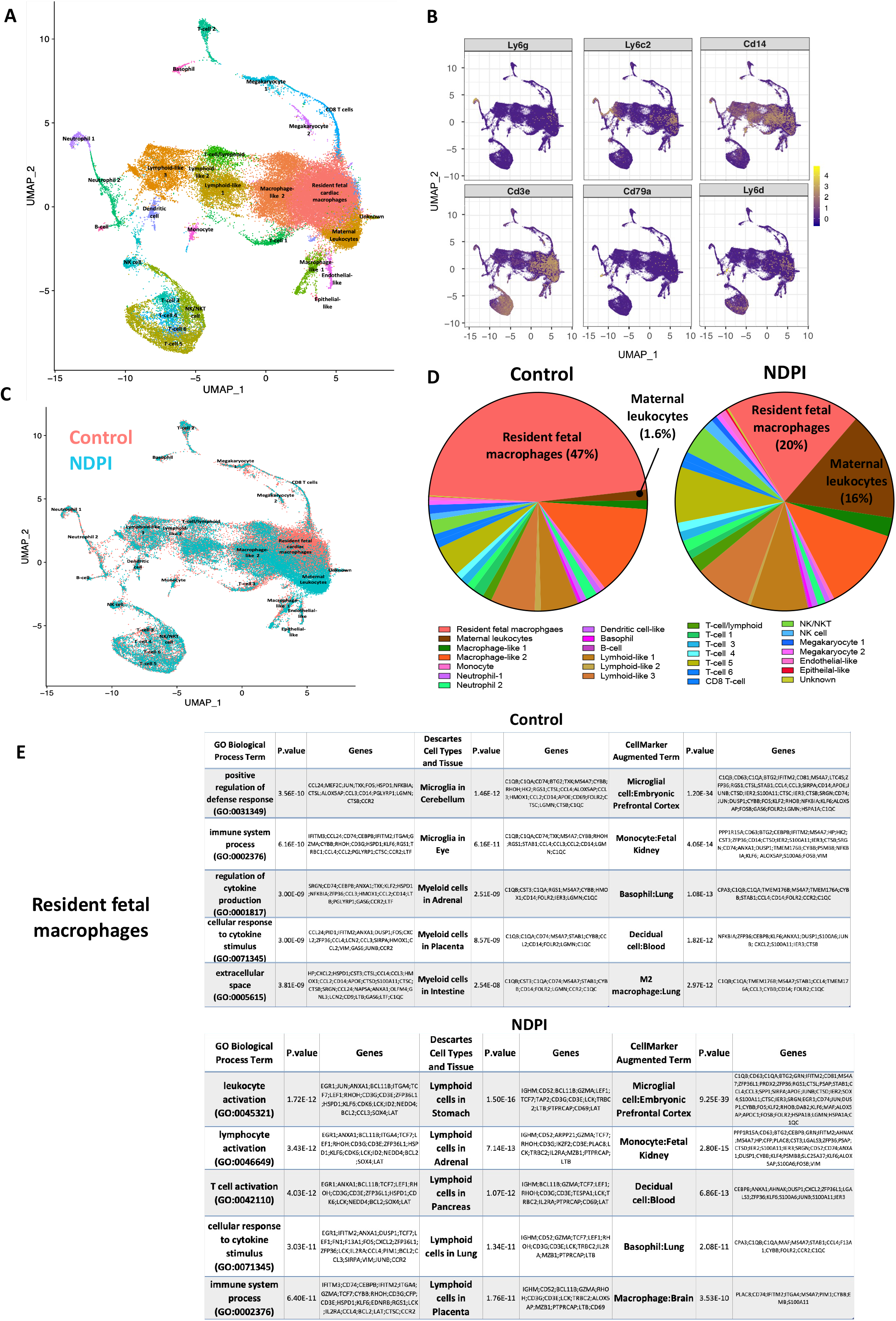
*Embryonic hearts from NDPI pregnancies have an altered* immune composition of. Neutrophils were depleted at day 4.5 and 7.5 of pregnancy using αLy6G. Mice were sacrificed at E14.5 of pregnancy and hearts dissected from harvested embryos from control or neutrophil depleted placental inflammation (NDPI) pregnancies. CD45^+^ cells were isolated from heart single cell suspensions using CD45 PE and anti-PE microbeads. Single cell sequencing was performed on the isolated cells. (A) Cells were clustered based on nearest neighbor clustering approach plotted by a UMAP. (B) UMAP showing cell clusters found in hearts of fetuses from control and NDPI pregnancies. (C) UMAP plots showing the level of expression of genes used to determine the different cell populations. (D) Comparison of the proportion of different leukocytes making up the total fetal heart leukocyte populations. (E) Tables showing the top 10 modulated GO biological process in the resident fetal macrophage populations in hearts from control or NDPI pregnancies.

### Impaired cardiac development from NDPI pregnancies persists into postnatal life

We next investigated whether the abnormal embryonic heart development detailed at E14.5 continues into postnatal life, assessed at postnatal day 5 (P5) and day 28 (P28).

Offspring from NDPI pregnancies at P5 presented with smaller body weight and a higher heart: body weight ratio, when compared to their control counterparts (Figure 6A). We observed a~90% reduction in maternal cells within P5 hearts (Figure 6B). In the resident CX_3_CR1^+^ leukocyte population, there was no significant difference in absolute numbers (Figure 6C), yet the proportion of CX_3_CR1^+^CCR2^−^ continued to be significantly lower in the hearts of offspring from NDPI pregnancies. The reverse occurred in offspring from control pregnancies (Figure 6D). As we observed hypertrabeculation of embryo hearts from NDPI pregnancies at E14.5, with a thinner LV wall (Figure 3), we assessed cardiac architecture of P5 hearts from both groups, staining cross-sections with DAPI, wheat germ agglutinin (for general architecture) and endomucin (endocardial cell marker). Cardiac tissue from P5 hearts from control pregnancies revealed a tight, compact structure, with normal endocardial formation as evidenced by high endomucin staining. In contrast, cardiac tissue of P5 hearts of offspring of NDPI pregnancies displayed hypertrabeculation, with a less compact endocardial structure coupled with a significant reduction in endomucin staining (Figure 6E) and was accompanied with a significant reduction in the number of CD31^+^ endothelial cells (Figure 6F).

**Figure 6:**
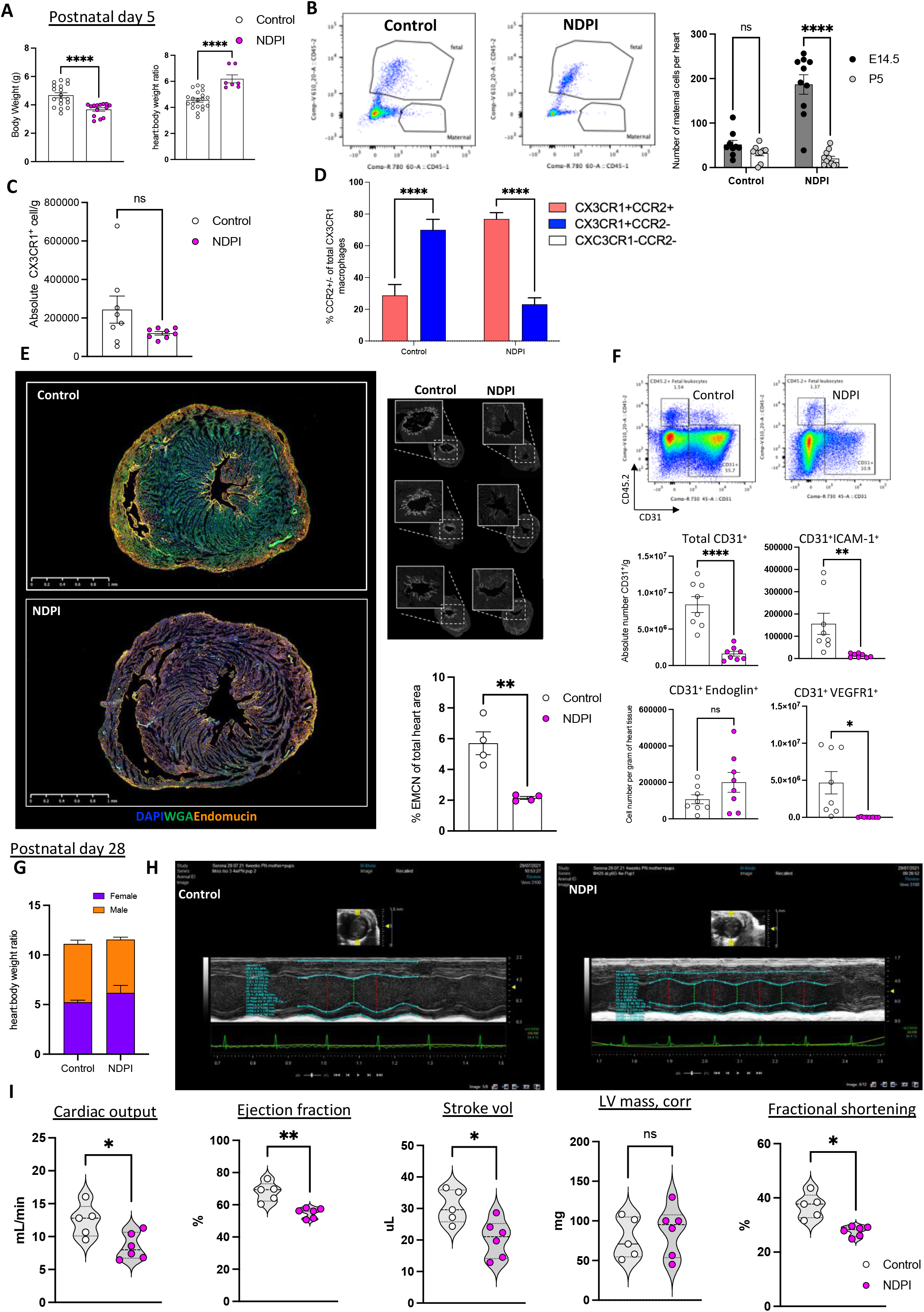
Aberrant embryonic cardiac development from NDPI pregnancies continues into postnatal life. Neutrophils were depleted at day 4.5 and 7.5 of pregnancy using αLy6G. Offspring of these dams were sacrificed at post-natal day 5 (P5) or 28 as indicated and heart structure and immune composition assessed. Offspring from Control (white) and neutrophil depleted (NDPI) (pink). (A) P5 offspring body weights in grams and heart: body weight ratio. (B) Flow cytometry plots of leukocytes in postnatal day 5 hearts from control of NDPI dams; CD45.1^+^ CD45.2^−^ cells are of maternal origin and CD45.1^+^CD45.2^+^ cells are of fetal origin. Graph showing number of maternal cells in offspring at E14.5 and P5. (C) Number of CX3CR1^+^ cells per P5 heart (D) Proportion of CCR2^+^ and CCR2^−^ cells within the total population of CX3CR1+ macrophages in P5 heart (E) Immunofluorescent images showing the expression of endomucin and wheat germ agglutinin in transverse sections of hearts from 4-week-old offspring from control or NDPI. Scale bar 1mm. Magnified images showing endomucin staining of the ventricle. Graph showing the proportion of total heart area expressing endomucin. (F) Representative FACs plots showing heart CD31^+^ populations from P5 offspring from dams treated with either isotype control or αLy6G neutralizing antibody. Graphs showing the number of CD31+ cells per gram of heart tissue and the number of these CD31^+^ cells expressing ICAM-1, Endoglin and VEGFR1. (G) Heart: body weight ratio of P28 male and female offspring from control or NDPI dams. (H) Representative echocardiography plots (I) Graphs showing quantification of heart parameters determined by echocardiography. Each symbol represents an individual mouse and statistical significance was tested by unpaired t-test. Ns = not significant, *p≤0.05, ** p≤0.01, ***p≤0.001 **** p≤0.0001. In all cases, data are mean ± SEM. MFI, median fluorescence intensity

We next analyzed hearts from adult offspring at P28. Measurements of body weights of offspring from both groups of pregnancies demonstrated no significant differences, suggesting post-weaning recovery by the NDPI offspring (Fig S5A). This result was matched by no difference in the heart: body weight ratios (Figure 6H). Phenotypic characterization of cardiac macrophages revealed the proportion of F4/80^+^Ly6C^+^ macrophages was significantly higher in NDPI offspring hearts, with augmented cell surface levels of MHC II and CCR2 and a reciprocal decrease in MertK expression (Figures S5B and S5C). We next sought to investigate the cardiac function in the offspring. Echocardiography of female offspring from NDPI pregnancies revealed a significant attenuation of cardiac output, stroke volume, ejection fraction and fractional shortening, with no difference in left ventricle mass, compared to control (Figure 6I and 6J). Taken together, these data suggest a functional continuity between cardiac abnormalities in utero at E14.5 in NDPI pregnancies with an impairment in cardiac function into postnatal and early adult life.

### Quelling placental inflammation prevents abnormal cardiac development

Based on our data, we propose a model where a breakdown in the placental tissue barrier, because of high local inflammation, promotes cardiac-selective influx of proinflammatory maternal leukocytes which impacts on normal cardiac development. In the final set of experiments, we challenged this hypothesis by dampening exaggerated NDPI expecting to restore normal fetal cardiac development. Thus, we targeted TNF-α-mediated placental inflammation by injecting NDPI pregnant mice with a neutralizing anti-TNF-α antibody (referred hereafter as NDPI+aTNF-α; see Figure S6A for scheme). Blocking TNF-α reduced placental TNF-α levels and i) rescued the poor placental tissue architecture, ii) enabled deeper trophoblast invasion and iii) restored placenta collagens at both gene and protein level (Figure S6B-E). Fewer activated placental neutrophils were counted, expressing lower levels of TNF-□, CXCR2, CD114 and MMP9 from NDPI+aTNF-α pregnancies compared to NDPI; in turn, this resulted in reduced number of inflammatory monocytes (Figure 7 A and B). Moreover, following maternal aTNF-α treatment, fewer maternal cells were recruited into the embryonic hearts (Figure 7C), with no difference in total fetal leukocyte number (data not shown). However, following maternal aTNF-α treatment there was up-regulation in the proportion of CX_3_CR1^+^CCR2^−^ resident fetal macrophages when compared to both control and NDPI pregnancies (Figure 7D) and a significant attenuation in the number of CX_3_CR1^+^CCR2^+^ cells per embryo heart, compared to NDPI alone (Figure S6F). This was coupled with a downregulation in both IL-1β and IL-6 gene expression. These cellular, molecular and histological changes yielded a well-defined cardiac structure, resembling the structure of control embryonic hearts including left ventricle wall thickness, coupled with a restoration in CD31^+^ endothelial cell numbers (Figure 7F and Fig S6G).

**Figure 7:**
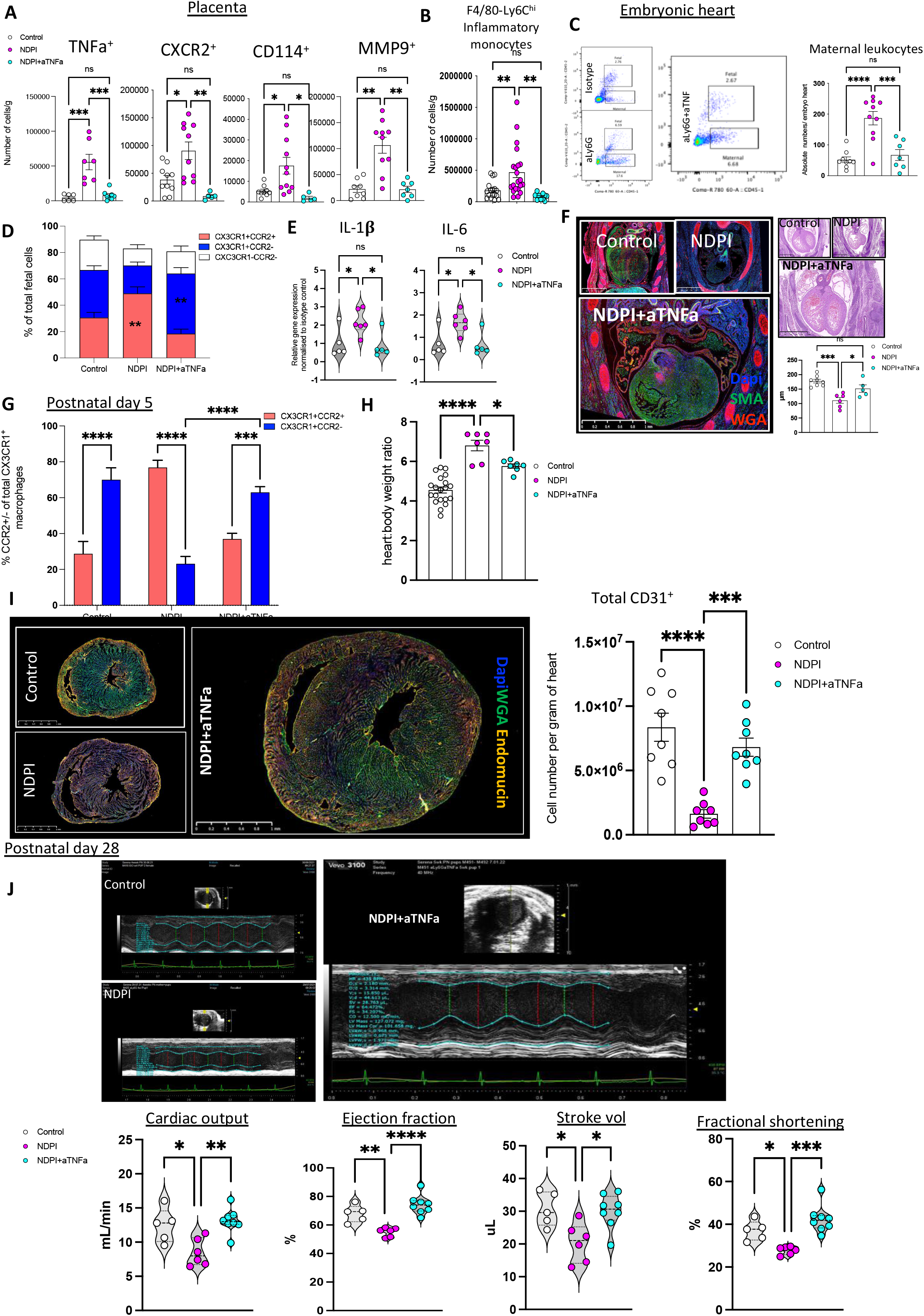
Quelling placental inflammation prevents abnormal cardiac development. Neutrophils were depleted at day 4.5 and 7.5 of pregnancy using αLy6G. At day 8.5 of pregnancy TNFα was neutralized by injecting αTNFα IgG i.v. Mice were sacrificed at E14.5 of pregnancy and placentae and embryos harvested to assess the structure and immune composition. Isotype control treated (white), neutrophil depleted (NDPI) (pink), neutrophil depleted TNFα neutralized (NDPI + aTNFa) (blue). (A) Absolute number of neutrophils expressing TNFα, CXCR2, CD114 and MMP9 from placentae analyzed by flow cytometry expressed as absolute cell number per gram of tissue. (B) Inflammatory monocytes from placentae analyzed by flow cytometry expressed as absolute cell number per gram of tissue. (C) Representative FACS plots showing fetal and maternal leukocytes in embryo heart and graph showing quantification of number of maternal leukocytes per embryo heart. (D) Proportion of total cells in embryo heart expressing CX3CR1^+^CCR2^+^, CX3CR1^+^CCR2^−^ or CX3CR1^−^CCR2^−^. (E) Violin plot showing gene expression in embryo hearts from control and NDPI pregnancies normalized to control. (F) Immunofluorescent images of E14.5 embryo heart of cross section. Sections were stained with DAPI (blue) WGA (red) SMA (green). Graph showing ventricle wall thickness quantification. (G) Proportion of CCR2^+^ and CCR2^−^ cells within the total population of CX3CR1+ macrophages in P5 heart. (H) Heart: body weight ratio of P5 offspring from control, NDPI, NDPI + aTNFa dams. (I) Immunofluorescent images showing the expression of endomucin (orange) and WGA (green) in cross sections of hearts from P5. Scale bar 1mm. Magnified images showing endomucin staining of the ventricle. Graph showing the proportion of total CD31^+^ cells in P5. (J) Echocardiography plot with graphs (bottom panel) showing quantification of heart parameters determined by echocardiography. Each symbol represents an individual mouse and statistical significance was tested by Oneway or Two-way ANOVA with Bonferroni post-correction. ns = not significant, *p≤0.05, ** p≤0.01, ***p≤0.001 **** p≤0.0001. In all cases, data are mean ± SEM. MFI, median fluorescence intensity

With respect to postnatal life, we observed a ~2-fold increase in the proportion of CX_3_CR1^+^CCR2^−^ macrophages relative to CX_3_CR1^+^CCR2^+^ cells, in NDPI+aTNF-α P5 hearts and compared to P5 hearts from NDPI offspring (Figure 7G). Heart: body weight ratios from offspring of NDPI+aTNFα were akin to control offspring (Figure 7H). When tissue architecture of P5 hearts was analyzed, there was a loss in the hypertrabeculation displayed by NDPI pregnancies, and a more compact endocardium within the cardiac tissue, accompanied by a significant increase in CD31^+^ endothelial cells (Figure 7I).

Finally, we assessed cardiac function of P28 hearts. Cardiac function was restored in NDPI+aTNF-α offspring, as cardiac output, stroke volume, ejection fraction and fractional shortening returned to control offspring levels (Figure 7J). This more normal cardiac functionality coincided with fewer F4/80^+^Ly6C^+^ macrophages within the cardiac tissue that displayed a more quiescent, anti-inflammatory cardiac macrophage phenotype, expressing significantly lower levels of MHC II and CCR2 and significantly higher levels of MertK compared to their NDPI offspring counterparts (Figure S6H).

Taken together these data suggest tempering neutrophil-driven placental inflammation with TNF-α neutralization is sufficient to rescue cardiac development and function in offspring postnatally and into adulthood.

## Discussion

We describe how placental inflammation driven by activated neutrophils promote aberrant embryonic heart development that impacts cardiac structure and downstream cardiac function in postnatal life. Specifically, locally produced neutrophil-derived TNF-□ promotes the establishment of an inflammatory placental environment with breakdown of tight tissue barriers; this, in turn, allows the transfer of placental inflammatory maternal monocytes to the embryonic heart, impeding its normal development, leading to manifest persistent heart dysfunction.

Following neutrophil depletion, an activated circulating neutrophil phenotype emerges, with high levels of CXCR2 and GCSF receptor CD114, and it is present within the placental tissue, coupled with high expression of TNF-α and MMP9. This neutrophil-driven inflammatory environment is responsible for matrix degradation of the placentae, with repressed collagen gene and protein expression.

The complex structure of the placenta barrier ensures maternal leukocytes do not enter the fetal compartment, and this guarantees the tolerogenic state of the placenta^29^. However, maternal cells, can cross the barrier and *via* the fetal circulation (called maternal microchimeric cells), enter fetal organs including liver, lung, pancreas and heart^30, 31^. A recent study identified maternal microchimeric cells in cardiac tissue of infants who died from neonatal lupus with heart block^32^, suggesting cells of maternal origin may impact on cardiac responses in offspring. Utilizing the CD45.1 (maternal) and CD45.2 (paternal) system, we identified a significant increase in the number of maternal (CD45.1^+^CD45.2^−^) inflammatory macrophages in embryonic hearts from NDPI pregnancies. The phenotype of the maternal cells is akin to the maternal inflammatory monocytes detected in the placenta. Analyses of chemokines in embryonic hearts and livers identified an embryonic heart-specific (but not liver) chemokine ligand profile resulting from NDPI pregnancies (namely CCL3, CCL4), which matched reciprocal chemokine receptors on maternal cells both in placenta and embryonic heart. This raises the question of what may be responsible for altering the environment of the developing heart? The timing of both neutrophil depletions in our model precedes maternal inflammatory immune cell infiltration since the 1^st^ depletion is carried out at E4.5, which is prior to both placenta and heart development, and the 2^nd^ depletion at E7.5, at the beginning of placental and heart development^33^. This suggests manipulating the maternal quiescent neutrophil response in early gestation may prime the placenta and - subsequently – the embryonic heart to a pro-inflammatory environment.

Placental development occurs in parallel to embryonic cardiac development and studies have highlighted the reciprocal influence these organs exert on each other’s development^1, 3, 34, 35^. Embryonic hearts from NDPI pregnancies display hypertrabeculation and thinner left ventricular walls, reminiscent of the congenital heart condition left ventricular non-compaction^23, 24^. Recent elegant studies have demonstrated that resident cardiac fetal macrophages populate distinct regions of the developing heart: while CX3CR1^+^CCR2^−^ macrophages predominantly populate the myocardial wall, CX3CR1^+^CCR2^+^ macrophages are found in the trabecular projections^12^, however the exact function during embryonic heart development of this latter subtype is currently unclear. Noteworthy, the presence of CCR2^+^ macrophages within the embryonic and neonatal heart is short lived, whereas their CCR2^−^ counterparts is self-renewing^13, 27^. The current data add another layer of complexity to this dichotomy, whereby presence of inflammatory maternal cells within the developing heart can promote the induction of these CCR2^+^ macrophages. In adult cardiac tissue, the roles of CCR2^−^ and CCR2^+^ macrophages are more apparent: resident CCR2^−^ macrophages of fetal origin have a reparative function, whereas CCR2^+^ monocyte-derived macrophages promote myocardial injury and inflammation^36^. Our findings indicate that, within the embryonic heart, maternal inflammatory cells can skew toward a fetal CCR2^+^ ‘inflammatory’ macrophage phenotype, that coincides with a distinct LVNC tissue architecture. These fetal CCR2^+^ macrophages may function to promote embryonic cardiac injury *or* they may create an inflammatory environment subsequently impacts heart development and may suggest why these cells are short-lived under normal heart developmental conditions. Postnatally, while there was no presence of inflammatory maternal cells in offspring hearts, an inflammatory cardiac macrophage phenotype persisted in P5 and P28 hearts from offspring of NDPI pregnancies, a cellular feature coupled with poor cardiac tissue architecture and function. Altogether, these data indicate that the influence of inflammatory maternal cells in embryonic heart development is imprinted in postnatal life and adds support to the concept that the innate immune competency of the placenta is key to the leukocyte composition of the developing heart.

In summary, we present a mechanistic paradigm whereby neutrophil-driven inflammation in pregnancy can preclude normal embryonic heart development as a direct consequence of poor placental development. Importantly, our study also opens translational avenues for early diagnoses and potential treatment of CHDs in utero. The former assertion is evidenced by our finding that NDPI pregnancies associate with an activated neutrophil phenotype within the maternal circulation. Relevantly, women with preeclampsia (PE) present an activated^16, 22^ and may produce a higher incidence of CHDs in offspring^37,38^.

Thus, early phenotyping of neutrophils from pregnant women could provide an early diagnostic tool to identify CHDs in fetuses. Finally, early identification of placental inflammation could enable therapeutic intervention like anti-TNF-α used here. Such an antiinflammatory therapy can restore normal embryonic and postnatal heart development and function, thus offering a potential alternative to current invasive in utero interventions to treat CHDs.

## Acknowledgments

This work was funded by a British Heart Foundation Intermediate Basic Science Research Fellowship (SN; FS/17/1/32528). The flow cytometry core facility was funded by CRUK (Core Award (C16420/A18066). We would like to thank Mr Reiss Brown and Mr Mason Arnold at the Biological Services Unit, Queen Mary, London, for their invaluable assistance with the animals during COVID lockdown.

## Author contributions

E.J.W carried out experiments, analyzed data and helped write the manuscript; S.B carried out experiments and analyzed data; S.F carried out experiments and analyzed data; N.P.D carried out experiments, analyzed data and helped write the manuscript; K.M.M and R.T.M analyzed RNA-seq data and provided figures; F.P. processed and produced HREM images; L.K.V and A.P.P provided human placenta tissue; M.P and F.M M-B provided critical discussion of experiments and the manuscript; S.N devised the concept, planned, carried out and supervised experiments; analyzed data, wrote the manuscript and composed figures.

## Resource availability

Raw bulk RNA-seq and scRNAseq data will be published in a relevant repository prior to publication. Further information and requests for resources and reagents should be directed to and will be fulfilled by the lead contact, Suchita Nadkarni.

## Methods

### Animals

Female C57/Bl6J mice and male Balb/C mice were purchased from Charles River UK. CD45.1 (B6.SJL-Ptprca Pepcb/BoyJ) were originally purchased from Charles River Italy and The Jackson Laboratory, respectively. All mice were housed at the animal facility of Queen Mary University of London and were conducted with strict adherence to the Home Office guidelines (PPL P71E91C8E) following approval by the Animal Use and Care Committee of Queen Mary University of London (QMUL). Mice used in the experiments of this study, unless specified were used at 8-12 weeks old.

### Human Samples

This study was carried out following the Declaration of Helsinki, was approved by the Research Ethics Committee of the University of Southern Santa Catarina (UNISUL) under 34681920.8.0000.5369 (for Preeclampsia samples - approved in 20/08/2020) or CAAE: 36084720.9.0000.5369 (for cardiac defects - approved in 18/01/2021) and all participants signed written informed consent and collected samples have been anonymised.

### Timed-mated allogeneic pregnancy

Timed mating was performed by housing Balb/C male mice with aged-matched female C57BL/6J or CD45.1 mice overnight and confirmed by the presence of a copulatory plug the following morning; this was defined as day 1 or mouse embryonic stage 0.5 (E0.5). Circulating maternal neutrophils were depleted at days 5 (E4.5) and 8 (E7.5) of pregnancy using a monoclonal neutralizing antibody (Biolegend, clone 1A8; 50 μg i.v.) or isotype control (Biolegend clone RTK2758; 50μg i.v). Following neutrophil depletion, some mice were injected with anti-mouse TNF-α i.v (Biolegend clone MP6-XT22; 10mg/kg i.v) or Progesterone (Sigma, 4-Pregnene-3,20-dione; 1mg s.c) on day 9 of pregnancy. Pregnant females were sacrificed at either day 15 of gestation (E14.5) or left to give birth and sacrificed at post-natal day 5 (P5) or post-natal day 28 (P28).

### Placental Neutrophil isolation

Placentas obtained from E14.5 pregnant mice were cut into small fragments and digested in PBS with 20mM Hepes, 60 U/ml DNase I and 450 U/ml Collagenase I at 37°C and 250rpm for 20 minutes. Suspensions were passed through 70μm filters, washed and red blood cells lysed with ACK lysis buffer. Placental neutrophils contained in the resulting single cell suspension were enriched via positive selection using 0.2μg anti-mouse Ly6G PE (Biolegend, clone 1A8) per 1×10^6^ cells followed by anti-PE microbeads (Biolegend), used according to the manufacturer’s instructions. The enriched placental neutrophil population were then used in downstream applications.

### Fetal heart CD45^+^ cell isolations

Hearts from fetuses at E14.5 were dissected and passed through a 70μm cell strainer to obtain a single cell suspension. Heart CD45^+^ cells were enriched via positive selection using 0.2μg anti-mouse CD45 PE (Biolegend, clone 30-F11) per 1×10^6^ cells followed by anti-PE microbeads (Biolegend), used according to the manufacturer’s instructions. The enriched fetal heart CD45^+^ cells were subsequently used for single cell RNA sequencing.

### Placental neutrophil splenic monocyte *in vitro* co-culture

Placental neutrophils were obtained as outlined above. Monocytes, from spleens of non-pregnant female C57BL/6J mice were isolated using a commercially available negative selection kit according to manufacturer’s instructions (EasySep mouse monocyte isolation kit). 1×10^5^ placental neutrophils were co-cultured with 1×10^5^ splenic monocytes in DMEM (Gibco) growth media containing 10% FCS and 50 IU/mL penicillin, 50 μg/mL streptomycin and supplemented with 10ng/ml M-CSF (Biolegend; cat#576402) at time of culture and again at day 3 of culture. Cells were cultured for a total of 5 days.

### Flow cytometry and antibodies

Single cell suspensions were prepared from uterine draining lumbar lymph nodes, nondraining cervical and spleen. Dead cells were excluded using a fixable viability dye LIVE/DEAD fixable aqua (Thermo Fisher Scientific, L34957). Cells were gated on singlets. Within the single cell population, dead cells were excluded using a fixable viability dye LIVE/DEAD fixable aqua (Thermo Fisher Scientific, L34957). Cells were then further phenotype as indicated in the results. Representative FACS plots show initial gating strategy for singlets and live cells are shown below.

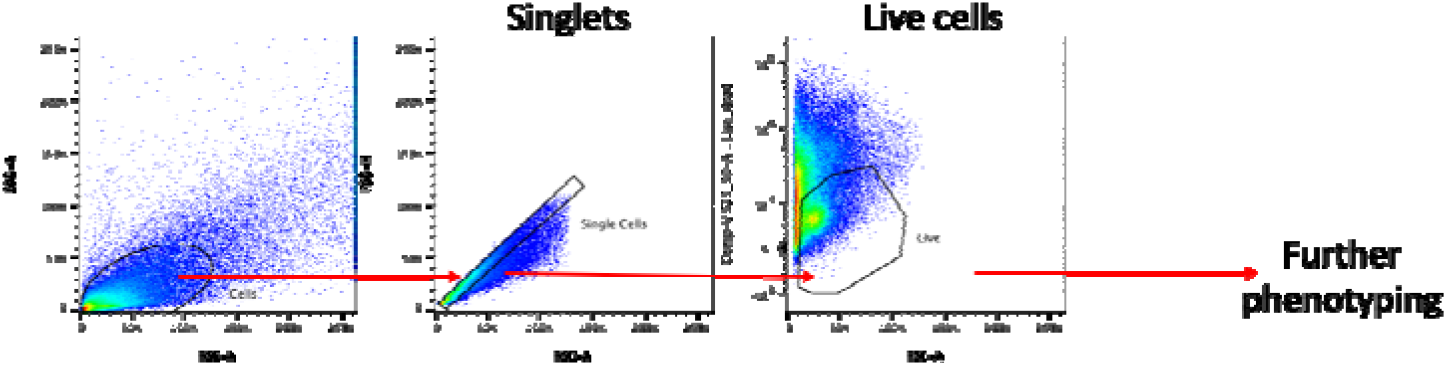

### Immunofluorescence microscopy

Tissues were fixed in 4% paraformaldehyde for 2□hours and paraffin or OCT embedded. Paraffin embedded sections were de-paraffinised with Histoclear and re-hydrated prior to antigen-retrieval performed by heating sections to 95°C for 10 min in Abcam antigen retrieval solution (ab64236). Sections were stained for hematoxylin/eosin or incubated with primary antibodies for immunofluorescence staining and mounted with Fluoromount G (Thermofisher 00-4958-02)

The primary antibodies used are listed in the table below. Nuclei were visualized using DAPI. Wide field imaging (bright field and immunofluorescence) was carried out using Nanozoomer S210 or S60 slide scanner (Hamamatsu). Confocal microscopy was carried out on a Carl Zeiss LSM800. Images were analysed with ImageJ (NIH) and Volocity (Version 6.3, PerkinElmer).

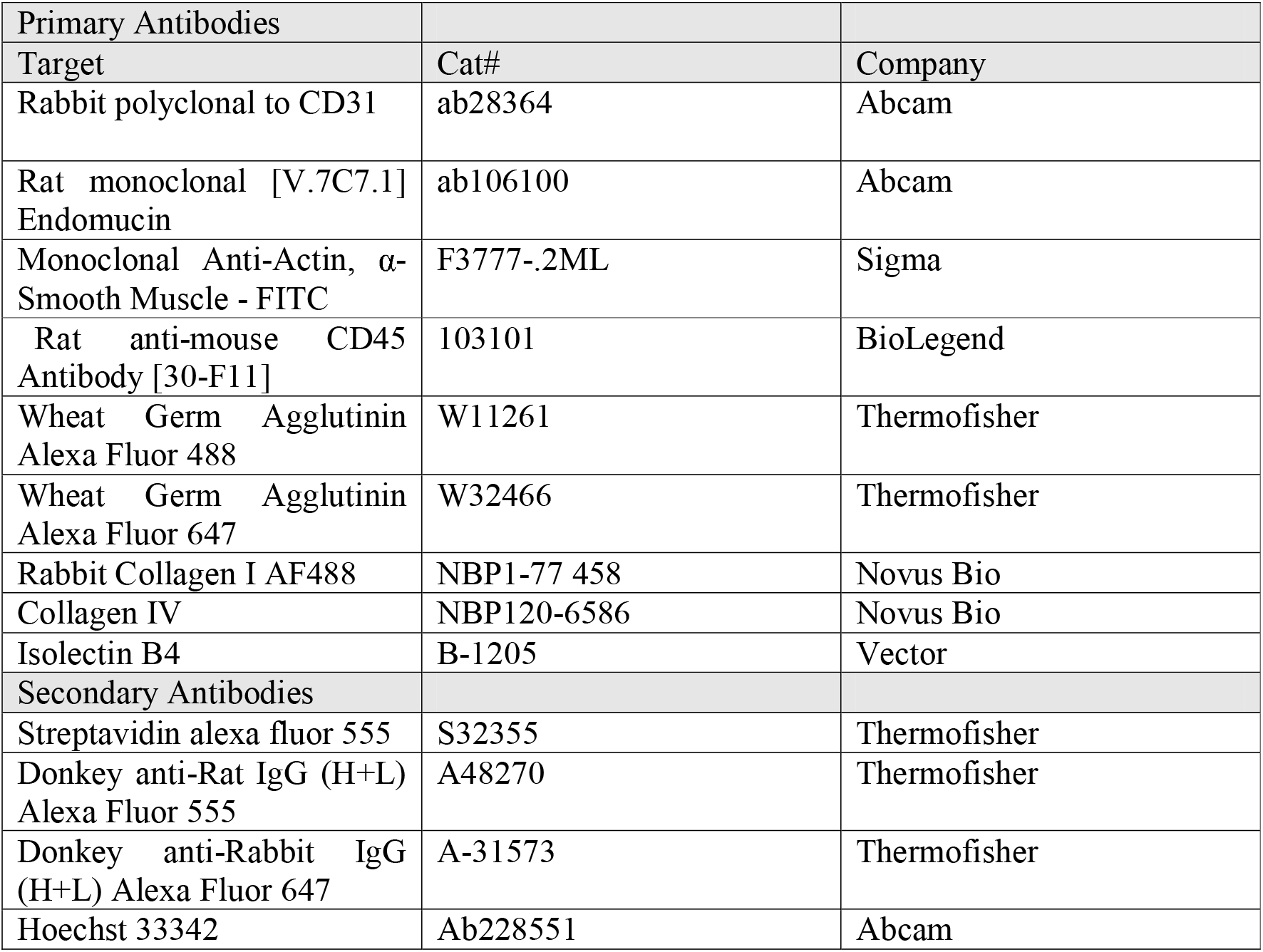

### High resolution episcopic microscopy (HREM)

Samples for high resolution episcopic microscopy (HREM) were fixed in Bouin’s for a minimum of 12h followed by extensive washing in PBS, dehydration in graded methanol series, incubation in JB-4 (Sigma) /Eosine (Sigma)/Acridine orange (Sigma) mix overnight to ensure proper sample infiltration and then embedded in fresh mix by adding the JB4 accelerator. Once polymerized the block where imaged as previously described^39, 40^. Samples were sectioned on a Leica sledge microtome at 1 or 2 μm or on commercial oHREM (Indigo Scientific) at 0.85 or 1.7 μm. An image of the surface of the block was then acquired under GFP excitation wavelength light using Olympus MVX10 microscope and High-resolution camera (Jenoptik). After acquisition the stacks were adjusted for gray level using Photoshop CS6 and then process for isotropic scaling, ortogonal re-sectioning, 25% downscaling, using a mixture of commercial and homemade software (see Wilson R et al. NAR 2016, Vol. 44 D855-D861). 3D volume rendering of the datasets were mostly produce from the 25% downscale stack using OsirixMD or Horos.

### Cytokine analysis

Placenta digest supernatants and plasma concentrations of TNF-α (Peprotech, 900-TN54) and IL-1β (Biolegend, 432601) were measured by ELISA according to the manufacturer’s instructions.

### *In Vivo* Cardiac Functional Assessments

Echocardiography was performed on mice to assess cardiac structure and function on day post-natal day 28 using a Vevo-3100 imagining system (VisualSonics, Toronto, Ontario, Canada). Echocardiography was performed on mice to assess cardiac structure and function at 4 weeks of age, using the Vevo-3100 imagining system (VisualSonics, Toronto, Ontario, Canada). Mice were anaesthetized (3% isoflurane in 0.2L/min oxygen, maintained with 2% isoflurane and 0.4L/min) and body temperature maintained at 37 °C. Heart rate was monitored for the duration of the procedure. Images from two-dimensional brightness mode (B-mode) and one-dimensional motion mode (M-mode), were obtained. Left ventricular ejection fraction (EF%), fractional shortening (FS%), cardiac output and stroke volume measurements were obtained from short-axis M-mode. Values were averaged from three beats. Echocardiographic analyses were performed by two independent operators, using Vevo LAB dongle software V3.2.6.

### Quantitative Real-time polymerase chain reaction

Placenta, fetal heart and fetal liver samples were stored in RNAlater (Invitrogen) at −80°C until processing. 30mg of placenta tissue was disrupted using a sample grinding kit (GE healthcare) and RLT buffer containing β-mercaptoethanol. Fetal heart and liver tissues were disrupted using a syringe and 25G needle and RLT buffer containing 2-Bme. RNA from the disrupted placental tissue and fetal heart and liver tissue was extracted using the RNeasy mini kit, according to the manufacturer’s recommendations. RNA quality and quantity was assessed using absorption measurements (Roche, Nanodrop) prior to performing the reverse transcription reaction, as per the manufacturer’s instruction (BioRad, iScript™ cDNA synthesis kit). qPCR was performed using 10ng cDNA, 1μM PCR primers (see table below for sequences) and 1x SYBR GreeN PCR mastermix (BioRad) in a CFX connect light cycler (BioRad). Expression was calculated according to the method outlined by Vandesompele *et al* ^41^whereby data is normalized to two housekeeping genes, HPRT and RPL32A. The thermal cycling conditions were 95°C for 30 seconds and 40 cycles of 95°C for 5 seconds followed by 60°C for 30 seconds and then melt curve analysis 95°C for 5 seconds, 65°C for 5 seconds and 95°C for 5 seconds. Experiments were performed in duplicate.

### Bulk RNA sequencing

High-quality RNA was extracted from 3 individual placenta from. Each treatment group, isotype (control) and aLy6G (NDPI) using Qiagen RNA extraction kit (Qiagen, Germany) and submitted to Eurofins (Germany) for Illumina sequencing. Please see the analysis pipeline at github.com/kerrimalone/aLy6G_RNAseq. Briefly, read quality was assessed with FastQC (v0.11.9)(http://www.bioinformatics). Transcript quantification was performed using Salmon (v.0.14.2) and the *Mus musculus* reference transcriptome (GRCm38, Gencode, release M23)^42^. Transcripts were summarized at the gene level, genes displaying expression levels below one count per million (CPM) in at least 3 individual libraries were filtered out. Differential expression (DE) analysis was performed between NDPI and control using EdgeR ^43^[3]. A False-Discovery Rate (FDR) threshold (≤0.05, Benjamini-Hochberg) was set to define DE genes. Positive log2 fold change indicates that a gene is higher expressed in aLyG6-treated samples and vice versa. Cellular functions and pathways over-represented for the DE gene list were assessed using the SIGORA in R^44^.

### Single cell sequencing

Fetal heart CD45 cells were fixed using 80% methanol and stored at −80°C according to a GENEWIZ-developed protocol. Upon shipping to the Genewiz facility cryopreserved cells were thawed, washed, and counted following the 10x Genomics protocol. Cell suspensions were loaded onto the 10x Genomics Chromium Controller targeting ~10,000 cells per sample for processing and microdroplet generation. The resulting libraries were sequenced with the Illumina^®^ HiSeq and resulted in 52,874 reads on average per cell, per sample. Primary data analyses were conducted in R v4.0.3 following the analysis strategy outlined https://www.ncbi.nlm.nih.gov/pmc/articles/PMC8238499/. Raw BC count matrices were imported to Seurat v4 with min.cells 3 and min.features 400. Prior to normalization, each sample was downsampled randomly to 50000 cells. Each sample was then normalised via the Log-Normalisation method (‘LogNormalize’) with scale.factor 10000. Prior to integration, 2000 variable features were identified using selection.method ‘vst’. For identifying integration anchors, 2000 anchor features were identified via the cca method - other parameters were idims 1:30, k.anchor 5, k.filter 200, and k.score 30. For the final integration, the following parameters were set: idims 1:30 and k.weight 100. Prior to downsteam analysis, cells with mitochondrial gene expression > 10% were filtered out, as were those with nCount_RNA > 40000 and nFeature_RNA > 5000. Clusters were identified in the integrated data using default parameters for FindNeighbors and FindClusters. Markers for each cluster were identified using FindMarkers with parameters logfc.threshold 0.0, test.use ‘LR’, min.pct 0.25, and min.diff.pct −Inf. Markers per cluster, per treatment were identified using FindAllMarkers and parameters logfc.threshold 0.0, test.use ‘LR’, min.pct 0.0, and min.diff.pct = −Inf. Volcano plots were generated via EnhancedVolcano. Gene Set Enrichment Analysis was performed via fgsea. Gene Ontology (GO) and KEGG enrichment was performed with topGO and KEGGprofile, respectively. Gene enrichment was also performed using enrichR. Further plots were generated via scDataviz.

## References

1. Burton, G.J. & Jauniaux, E. Development of the Human Placenta and Fetal Heart: Synergic or Independent? Front Physiol 9, 373 (2018).

2. Krishnan, A. et al. A detailed comparison of mouse and human cardiac development. Pediatr Res 76, 500–507 (2014).

3. Courtney, J.A., Cnota, J.F. & Jones, H.N. The Role of Abnormal Placentation in Congenital Heart Disease; Cause, Correlate, or Consequence? Front Physiol 9, 1045 (2018).

4. Radhakrishna, U. et al. Placental epigenetics for evaluation of fetal congenital heart defects: Ventricular Septal Defect (VSD). PloS one 14, e0200229 (2019).

5. Adams, R.H. et al. Essential role of p38alpha MAP kinase in placental but not embryonic cardiovascular development. Mol Cell 6, 109–116 (2000).

6. Barak, Y. et al. PPAR gamma is required for placental, cardiac, and adipose tissue development. Mol Cell 4, 585–595 (1999).

7. (NCARDRS), N.C.A.a.R.D.R.S. NCARDRS congenital anomaly statistics 2017. PHE publications gateway number GW-473 (2017).

8. Epelman, S., Lavine, K.J. & Randolph, G.J. Origin and functions of tissue macrophages. Immunity 41, 21–35 (2014).

9. Mass, E. et al. Specification of tissue-resident macrophages during organogenesis. Science 353 (2016).

10. Schulz, C. et al. A lineage of myeloid cells independent of Myb and hematopoietic stem cells. Science 336, 86–90 (2012).

11. Stremmel, C. et al. Yolk sac macrophage progenitors traffic to the embryo during defined stages of development. Nature communications 9, 75 (2018).

12. Leid, J. et al. Primitive Embryonic Macrophages are Required for Coronary Development and Maturation. Circ Res 118, 1498–1511 (2016).

13. Lavine, K.J. et al. Distinct macrophage lineages contribute to disparate patterns of cardiac recovery and remodeling in the neonatal and adult heart. Proceedings of the National Academy of Sciences of the United States of America 111, 16029–16034 (2014).

14. Bajpai, G. et al. Tissue Resident CCR2- and CCR2+ Cardiac Macrophages Differentially Orchestrate Monocyte Recruitment and Fate Specification Following Myocardial Injury. Circ Res 124, 263–278 (2019).

15. Epelman, S. et al. Embryonic and adult-derived resident cardiac macrophages are maintained through distinct mechanisms at steady state and during inflammation. Immunity 40, 91–104 (2014).

16. Nadkarni, S. et al. Neutrophils induce proangiogenic T cells with a regulatory phenotype in pregnancy. Proceedings of the National Academy of Sciences of the United States of America 113, E8415–e8424 (2016).

17. Daley, J.M., Thomay, A.A., Connolly, M.D., Reichner, J.S. & Albina, J.E. Use of Ly6G-specific monoclonal antibody to deplete neutrophils in mice. J Leukoc Biol 83, 64–70 (2008).

18. Malak, T.M. et al. Confocal immunofluorescence localization of collagen types I, III, IV, V and VI and their ultrastructural organization in term human fetal membranes. Placenta 14, 385–406 (1993).

19. Hudson, D.M. & Eyre, D.R. Collagen prolyl 3-hydroxylation: a major role for a minor post-translational modification? Connect Tissue Res 54, 245–251 (2013).

20. Takahara, K. et al. Type I procollagen COOH-terminal proteinase enhancer protein: identification, primary structure, and chromosomal localization of the cognate human gene (PCOLCE). J Biol Chem 269, 26280–26285 (1994).

21. Oefner, C.M. et al. Collagen type IV at the fetal-maternal interface. Placenta 36, 59–68 (2015).

22. Hu, Y. et al. Increased Neutrophil Activation and Plasma DNA Levels in Patients with Pre-Eclampsia. Thromb Haemost 118, 2064–2073 (2018).

23. Rhee, S. et al. Endothelial deletion of Ino80 disrupts coronary angiogenesis and causes congenital heart disease. Nature communications 9, 368 (2018).

24. Zhang, W., Chen, H., Qu, X., Chang, C.P. & Shou, W. Molecular mechanism of ventricular trabeculation/compaction and the pathogenesis of the left ventricular noncompaction cardiomyopathy (LVNC). Am J Med Genet C Semin Med Genet 163C, 144–156 (2013).

25. Rhee, S. et al. Endocardial/endothelial angiocrines regulate cardiomyocyte development and maturation and induce features of ventricular non-compaction. Eur Heart J 42, 4264–4276 (2021).

26. Sandireddy, R. et al. Semaphorin 3E/PlexinD1 signaling is required for cardiac ventricular compaction. JCI Insight 4 (2019).

27. Dick, S.A. et al. Three tissue resident macrophage subsets coexist across organs with conserved origins and life cycles. Sci Immunol 7, eabf7777 (2022).

28. Chen, E.Y. et al. Enrichr: interactive and collaborative HTML5 gene list enrichment analysis tool. BMC Bioinformatics 14, 128 (2013).

29. Ander, S.E., Diamond, M.S. & Coyne, C.B. Immune responses at the maternal-fetal interface. Sci Immunol 4 (2019).

30. Jonsson, A.M., Uzunel, M., Gotherstrom, C., Papadogiannakis, N. & Westgren, M. Maternal microchimerism in human fetal tissues. American journal of obstetrics and gynecology 198, 325 e321–326 (2008).

31. Nelson, J.L. The otherness of self: microchimerism in health and disease. Trends in immunology 33, 421–427 (2012).

32. Stevens, A.M., Hermes, H.M., Rutledge, J.C., Buyon, J.P. & Nelson, J.L. Myocardial-tissue-specific phenotype of maternal microchimerism in neonatal lupus congenital heart block. Lancet 362, 1617–1623 (2003).

33. Sones, J.L. & Davisson, R.L. Preeclampsia, of mice and women. Physiological genomics 48, 565–572 (2016).

34. Courtney, J. et al. Abnormalities of placental development and function are associated with the different fetal growth patterns of hypoplastic left heart syndrome and transposition of the great arteries. Placenta 101, 57–65 (2020).

35. Wilson, R.L., Troja, W., Courtney, J., Williams, A. & Jones, H.N. Placental and fetal characteristics of the Ohia mouse line recapitulate outcomes in human hypoplastic left heart syndrome. Placenta 117, 131–138 (2022).

36. Dick, S.A. et al. Publisher Correction: Self-renewing resident cardiac macrophages limit adverse remodeling following myocardial infarction. Nat Immunol 20, 664 (2019).

37. Liu, J. et al. There is a Strong Association between Early Preeclampsia and Congenital Heart Defects: A Large Population-Based, Retrospective Study. Gynecol Obstet Invest 86, 40–47 (2021).

38. Yilgwan, C.S. et al. Profile of congenital heart disease in infants born following exposure to preeclampsia. Plo Sone 15, e0229987 (2020).

39. Mohun, T.J. & Weninger, W.J. Imaging heart development using high-resolution episcopic microscopy. Curr Opin Genet Dev 21, 573–578 (2011).

40. Weninger, W.J. et al. Visualising the Cardiovascular System of Embryos of Biomedical Model Organisms with High Resolution Episcopic Microscopy (HREM). J Cardiovasc Dev Dis 5 (2018).

41. Vandesompele, J. et al. Accurate normalization of real-time quantitative RT-PCR data by geometric averaging of multiple internal control genes. Genome Biol 3, RESEARCH0034 (2002).

42. Patro, R., Duggal, G., Love, M.I., Irizarry, R.A. & Kingsford, C. Salmon provides fast and bias-aware quantification of transcript expression. Nat Methods 14, 417–419 (2017).

43. Robinson, M.D., McCarthy, D.J. & Smyth, G.K. edgeR: a Bioconductor package for differential expression analysis of digital gene expression data. Bioinformatics 26, 139–140 (2010).

44. Foroushani, A.B., Brinkman, F.S. & Lynn, D.J. Pathway-GPS and SIGORA: identifying relevant pathways based on the over-representation of their gene-pair signatures. PeerJ 1, e229 (2013).

