## Supplemental Figures for "Placental inflammation leads to abnormal embryonic heart development"

Figure S1

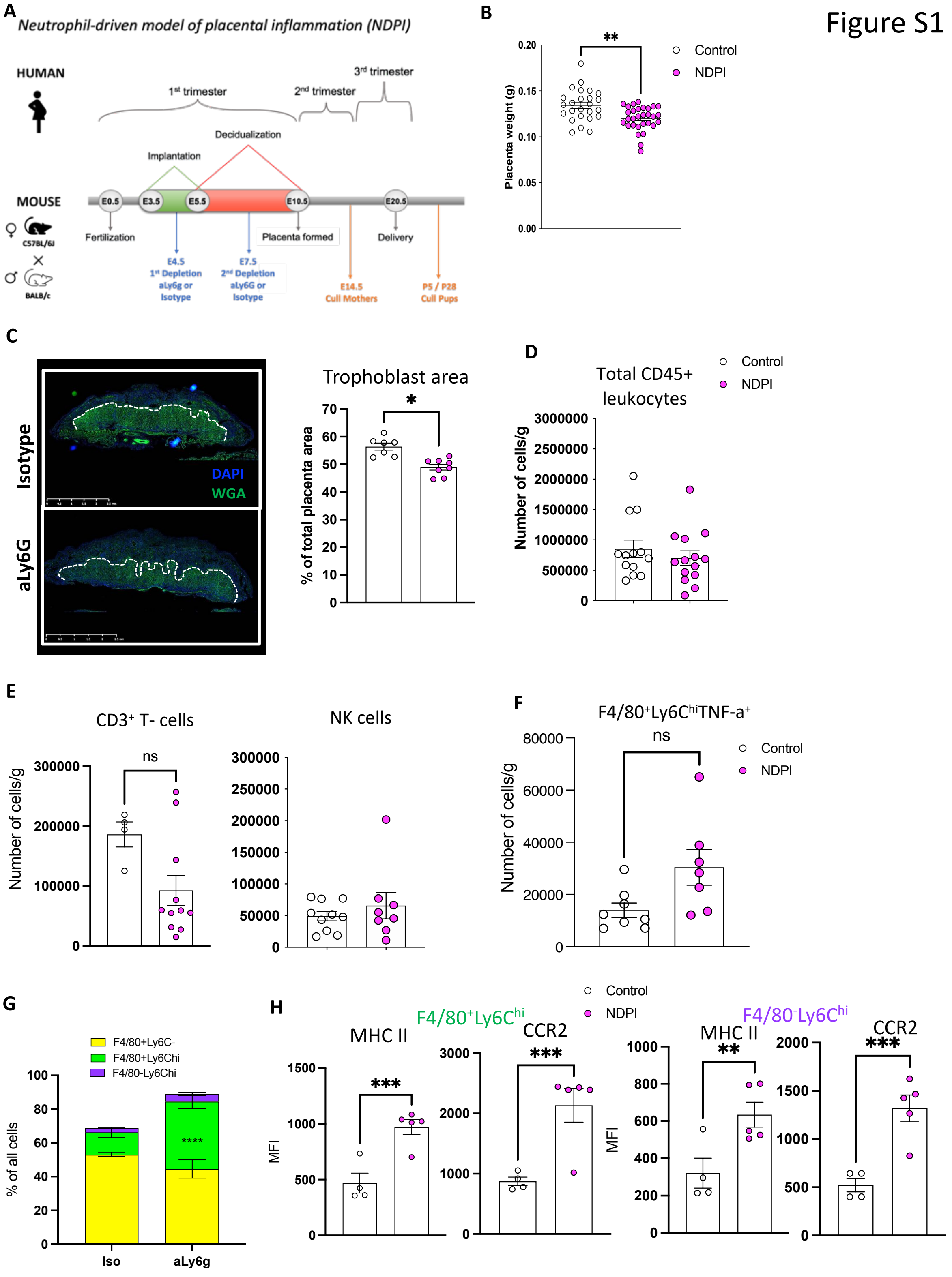

A
Neutrophil-driven model of placental inflammation (NDPI)
HUMAN
1st trimester
2nd trimester
3rd trimester
Decidualization
Implantation
MOUSE
C57BL/6J
BALB/c
E0.5
E3.5
E5.5
E10.5
E20.5
Fertilization
Placenta formed
Delivery
E4.5
1st Depletion
aLy6g or Isotype
E7.5
2nd Depletion
aLy6G or Isotype
E14.5
Cull Mothers
P5 / P28
Cull Pups
B
Placenta weight (g)
Control
NDPI
**
C
Isotype
aLy6G
DAPI
WGA
Trophoblast area
% of total placenta area
*
D
Total CD45+ leukocytes
Number of cells/g
Control
NDPI
E
CD3+ T- cells
ns
Number of cells/g
NK cells
F
F4/80+Ly6ChiTNF-a+
ns
Number of cells/g
Control
NDPI
G
F4/80+Ly6C-
F4/80+Ly6Chi
F4/80-Ly6Chi
% of all cells
****
Iso
aLy6g
H
Control
NDPI
F4/80+Ly6Chi
MHC II
CCR2
***
***
MFI
F4/80-Ly6Chi
MHC II
CCR2
**
***
MFI

***Supplementary figure 1***

Isotype control treated (white) and neutrophil depleted (NDPI) (pink).

(A) Schematic for experimental design

(B) Weights of E14.5 placentae.

(C) Immunofluorescent staining of placentae for wheat germ agglutinin (WGA) (green) to show gross structure and cell nuclei with DAPI (blue). White dotted line indicates the separation of the labyrinthine zone/trophoblast area and the outer junctional zone. Graph showing the proportion of the total area of the placentae occupied by the trophoblast zone.

(D-F) Flow cytometry quantification of different leukocyte subsets in placenta, expressed as cell number per gram of tissue (D) $CD45^+$ cells (E) $CD3^+$ total T cells, $NK1.1^+$ NK cells, (left to right) (F) $F4/80^+$ $Ly6c^{hi}TNF\text{-}\alpha^+$ macrophages.

(G) Proportion of $F4/80^+$ $Ly6C^-$, $F4/80^+Ly6C^{hi}$ and $F4/80^-Ly6C^{hi}$ cells making up the total leukocyte population in the placenta.

(H) $F4/80^+Ly6C^{hi}$ and $F4/80^-Ly6C^{hi}$ populations from placentae were analysed for the expression of MHCII and CCR2, expressed as median fluorescent intensity.

Each symbol represents an individual mouse and statistical significance was tested by unpaired t-test. ns = not significant, $*p\leq0.05$, $**\ p\leq0.01$, $***p\leq0.001$ $****\ p\leq0.0001$. In all cases, data are mean ± SEM. MFI, median fluorescence intensity

A

Figure S2

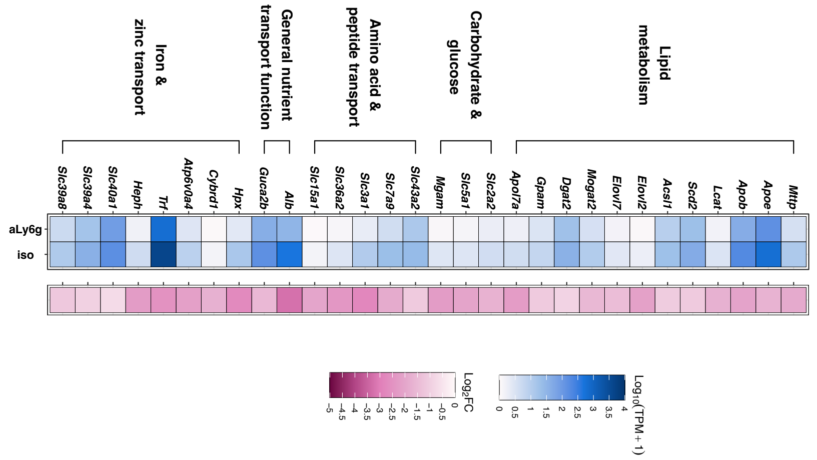


B

**Table 1 showing all 44 mouse collagens and their expression following RNA-seq analyses**

| Ensembl ID | Gene Symbol | Gene Name | Collagen Type | Adj P val | Log2FC |
|---|---|---|---|---|---|
| ENSMUSG00000001506 | Col1a1 | collagen, type I, alpha 1 (Col1a1), mRNA | type I | 3.27E-14 | 3.188074 |
| ENSMUSG00000029661 | Col1a2 | collagen, type I, alpha 2 (Col1a2), mRNA | type I | 3.45E-13 | 2.874913 |
| ENSMUSG00000022483 | Col2a1 | collagen, type II, alpha 1 (Col2a1), mRNA | type II | 3.6E-13 | 4.937598 |
| ENSMUSG00000026043 | Col3a1 | collagen, type III, alpha 1 (Col3a1), mRNA | type III | 0.442993343 | 0.681686 |
| ENSMUSG00000031502 | Col4a1 | collagen, type IV, alpha 1 (Col4a1), mRNA | type IV | 0.995426174 | 0.090912 |
| ENSMUSG00000031503 | Col4a2 | collagen, type IV, alpha 2 (Col4a2), mRNA | type IV | 0.995426174 | 0.103611 |
| ENSMUSG00000079465 | Col4a3 | collagen, type IV, alpha 3 (Col4a3), mRNA | type IV | 0.996574685 | 0.014233 |
| ENSMUSG00000067158 | Col4a4 | collagen, type IV, alpha 4 (Col4a4), mRNA | type IV | 0.989220301 | 0.464417 |
| ENSMUSG00000031274 | Col4a5 | collagen, type IV, alpha 5 (Col4a5), mRNA | type IV | 3.12E-14 | 3.130044 |
| ENSMUSG00000031273 | Col4a6 | collagen, type IV, alpha 6 (Col4a6), mRNA | type IV | 0.00000051 | 3.663765 |
| ENSMUSG00000026837 | Col5a1 | collagen, type V, alpha 1 (Col5a1), mRNA | type V | 0.000273973 | 1.73128 |
| ENSMUSG00000026042 | Col5a2 | collagen, type V, alpha 2 (Col5a2), mRNA | type V | 0.442105017 | 0.828681 |
| ENSMUSG00000004098 | Col5a3 | collagen, type V, alpha 3 (Col5a3), mRNA | type V | 0.989220301 | -0.32152 |
| ENSMUSG00000001119 | Col6a1 | collagen, type VI, alpha 1 (Col6a1), mRNA | type VI | 0.000000256 | 1.907141 |
| ENSMUSG00000020241 | Col6a2 | collagen, type VI, alpha 2 (Col6a2), mRNA | type VI | 0.0000231 | 1.60699 |
| ENSMUSG00000048126 | Col6a3 | collagen, type VI, alpha 3 (Col6a3), mRNA | type VI | 0.003193055 | 1.289803 |
| ENSMUSG00000032572 | Col6a4 | collagen, type VI, alpha 4 (Col6a4), mRNA | type VI | low expression | |
| ENSMUSG00000091345 | Col6a5 | collagen, type VI, alpha 5 (Col6a5), mRNA | type VI | 0.989220301 | 0.322323 |
| ENSMUSG00000043719 | Col6a6 | collagen, type VI, alpha 6 (Col6a6), mRNA | type VI | low expression | |
| ENSMUSG00000025650 | Col7a1 | collagen, type VII, alpha 1 (Col7a1), mRNA | type VII | 0.989220301 | -0.26505 |
| ENSMUSG00000068196 | Col8a1 | collagen, type VIII, alpha 1 (Col8a1), mRNA | type VIII | low expression | |
| ENSMUSG00000056174 | Col8a2 | collagen, type VIII, alpha 2 (Col8a2), mRNA | type VIII | 0.446143852 | 0.970351 |
| ENSMUSG00000026147 | Col9a1 | collagen, type IX, alpha 1 (Col9a1), mRNA | type IX | low expression | |
| ENSMUSG00000028626 | Col9a2 | collagen, type IX, alpha 2 (Col9a2), mRNA | type IX | 0.00000051 | 3.616545 |
| ENSMUSG00000027570 | Col9a3 | collagen, type IX, alpha 3 (Col9a3), mRNA | type IX | 0.000000014 | 3.965448 |
| ENSMUSG00000039462 | Col10a1 | collagen, type X, alpha 1 (Col10a1), mRNA | type X | low expression | |
| ENSMUSG00000027966 | Col11a1 | collagen, type XI, alpha 1 (Col11a1), mRNA | type XI | 1.04E-15 | 4.296395 |
| ENSMUSG00000024330 | Col11a2 | collagen, type XI, alpha 2 (Col11a2), mRNA | type XI | 0.00000051 | 2.899813 |
| ENSMUSG00000032332 | Col12a1 | collagen, type XII, alpha 1 (Col12a1), mRNA | type XII | 0.056573025 | 1.102192 |
| ENSMUSG00000058806 | Col13a1 | collagen, type XIII, alpha 1 (Col13a1), mRNA | type XIII | 0.996574685 | 0.015684 |
| ENSMUSG00000022371 | Col14a1 | collagen, type XIV, alpha 1 (Col14a1), mRNA | type XIV | 0.000000146 | 2.296451 |
| ENSMUSG00000028339 | Col15a1 | collagen, type XV, alpha 1 (Col15a1), mRNA | type XV | 0.989220301 | -0.18883 |
| ENSMUSG00000040690 | Col16a1 | collagen, type XVI, alpha 1 (Col16a1), mRNA | type XVI | 0.000372668 | 1.634199 |
| ENSMUSG00000025064 | Col17a1 | collagen, type XVII, alpha 1 (Col17a1), mRNA | type XVII | low expression | |
| ENSMUSG00000001435 | Col18a1 | collagen, type XVIII, alpha 1 (Col18a1), mRNA | type XVIII | 0.00000182 | 1.561112 |
| ENSMUSG00000026141 | Col19a1 | collagen, type XIX, alpha 1 (Col19a1), mRNA | type XIX | low expression | |
| ENSMUSG00000016356 | Col20a1 | collagen, type XX, alpha1 (Col20a1), mRNA | type XX | 0.99369326 | -0.12657 |
| ENSMUSG00000079022 | Col22a1 | collagen, type XXII, alpha 1 (Col22a1), mRNA | type XXII | 0.989220301 | -0.25757 |
| ENSMUSG00000063564 | Col23a1 | collagen, type XXIII, alpha 1 (Col23a1), mRNA | type XXIII | 0.995426174 | 0.053846 |
| ENSMUSG00000028197 | Col24a1 | collagen, type XXIV, alpha 1 (Col24a1), mRNA | type XXIV | low expression | |
| ENSMUSG00000058897 | Col25a1 | collagen, type XXV, alpha 1 (Col25a1), mRNA | type XXV | low expression | |
| ENSMUSG00000004415 | Col26a1 | collagen, type XXVI, alpha 1 (Col26a1), mRNA | type XXVI | low expression | |
| ENSMUSG00000045672 | Col27a1 | collagen, type XXVII, alpha 1 (Col27a1), mRNA | type XXVII | 0.010482257 | 1.417136 |
| ENSMUSG00000068794 | Col28a1 | collagen, type XXVIII, alpha 1 (Col28a1), mRNA | type XXVIII | low expression | |

***Supplementary Figure 2***

Experiment as outlined in S1-A. Mice were sacrificed at E14.5 of pregnancy and placentae harvested and processed for RNA sequencing.

(A) Heat map showing gene expression of nutrient transport pathways

(B) Gene expression of the 44 collagen genes of the mouse genome NDPI vs control.

A

Figure S3

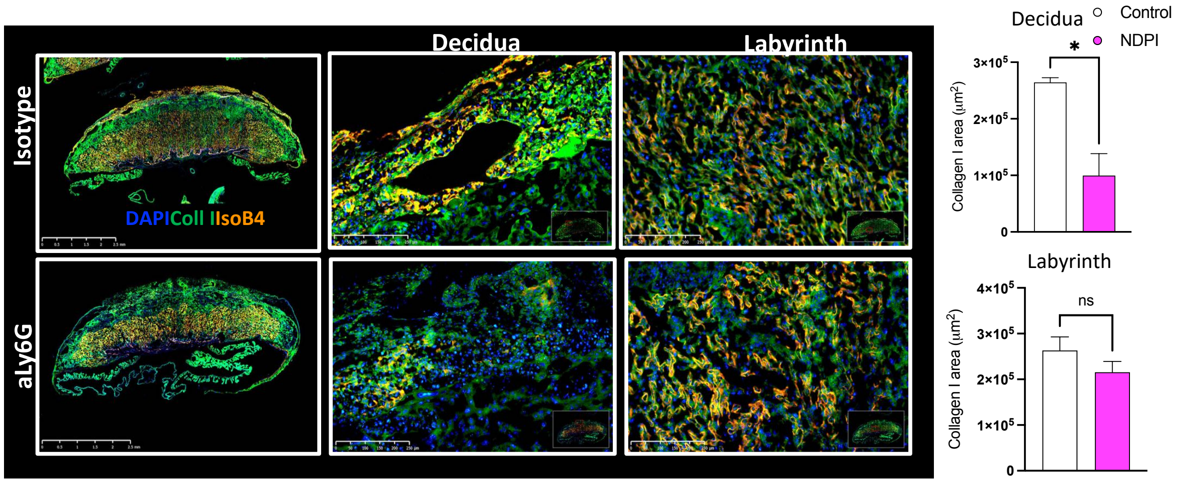


***Supplementary figure 3***

Experiment as outlined in S1-A. Mice were sacrificed at E14.5 of pregnancy and placentae harvested.

(A) Immunofluorescent staining of placentae for Collagen 1 (green) isolectin b4 (orange) and cell nuclei with DAPI (blue). Expression is shown in the whole placenta (left) decidual layer (middle) and labyrinth layer (right). Graphs showing quantification of the total area of Col I expression in the decidua and labyrinth.

Each symbol represents an individual mouse and statistical significance was tested by unpaired t-test. ns = not significant, *p≤0.05. Data are mean ± SEM.

Figure S4

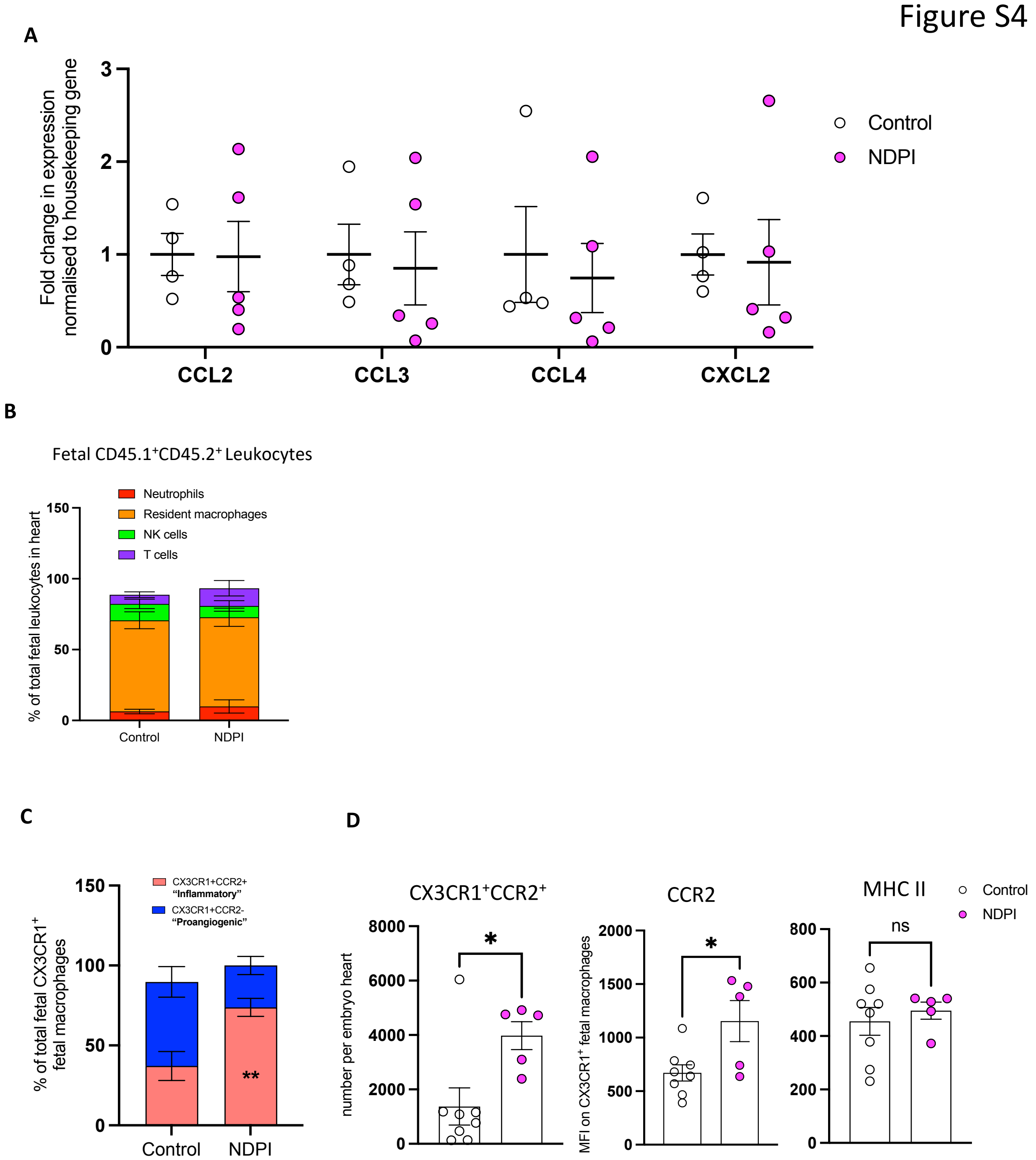

A
Fold change in expression normalised to housekeeping gene
0
1
2
3
CCL2
CCL3
CCL4
CXCL2
Control
NDPI
B
Fetal CD45.1+CD45.2+ Leukocytes
% of total fetal leukocytes in heart
Neutrophils
Resident macrophages
NK cells
T cells
0
50
100
150
Control
NDPI
C
% of total fetal CX3CR1+ fetal macrophages
CX3CR1+CCR2+ "Inflammatory"
CX3CR1+CCR2- "Proangiogenic"
0
50
100
150
**
Control
NDPI
D
CX3CR1+CCR2+
number per embryo heart
0
2000
4000
6000
8000
*
CCR2
MFI on CX3CR1+ fetal macrophages
0
500
1000
1500
2000
*
MHC II
ns
0
200
400
600
800
Control
NDPI

***Supplementary figure 4***

Neutrophils are depleted at day 4.5 and 7.5 of pregnancy using aLy6G. Mice are sacrificed at E14.5 of pregnancy and hearts dissected from harvested embryos. Isotype control treated (white) and neutrophil depleted (NDPl) (pink).

(A) Graph showing gene expression of CCL2, CCL3, CCL4 and CXCL2 in embryo livers from neutrophil depleted pregnancies normalised to isotype control.
(B) Proportion of fetal neutrophils, resident macrophages, NK (Natural Killer) cells and T cells making up the total fetal leukocyte populations in the fetal heart.
(C) Proportion of $CX3CR1^{+}CCR2^{+}$ antiangiogenic, $CX3CR1^{+}CCR2^{-}$ proangiogenic and $CX3CR1^{-}CCR2^{-}$ cells per embryo heart.
(D) Quantification of fetal heart antiangiogenic $CX3CR1^{+}CCR2^{+}$ cells. MFI of CCR2 and MHCII expression on $CX3CR1^{+}$ fetal macrophages.

Each symbol represents an individual mouse and statistical significance was tested by unpaired t-test. ns = not significant, *p≤0.05, ** p≤0.01. In all cases, data are mean ± SEM. MFI, median fluorescence intensity.

Figure S5

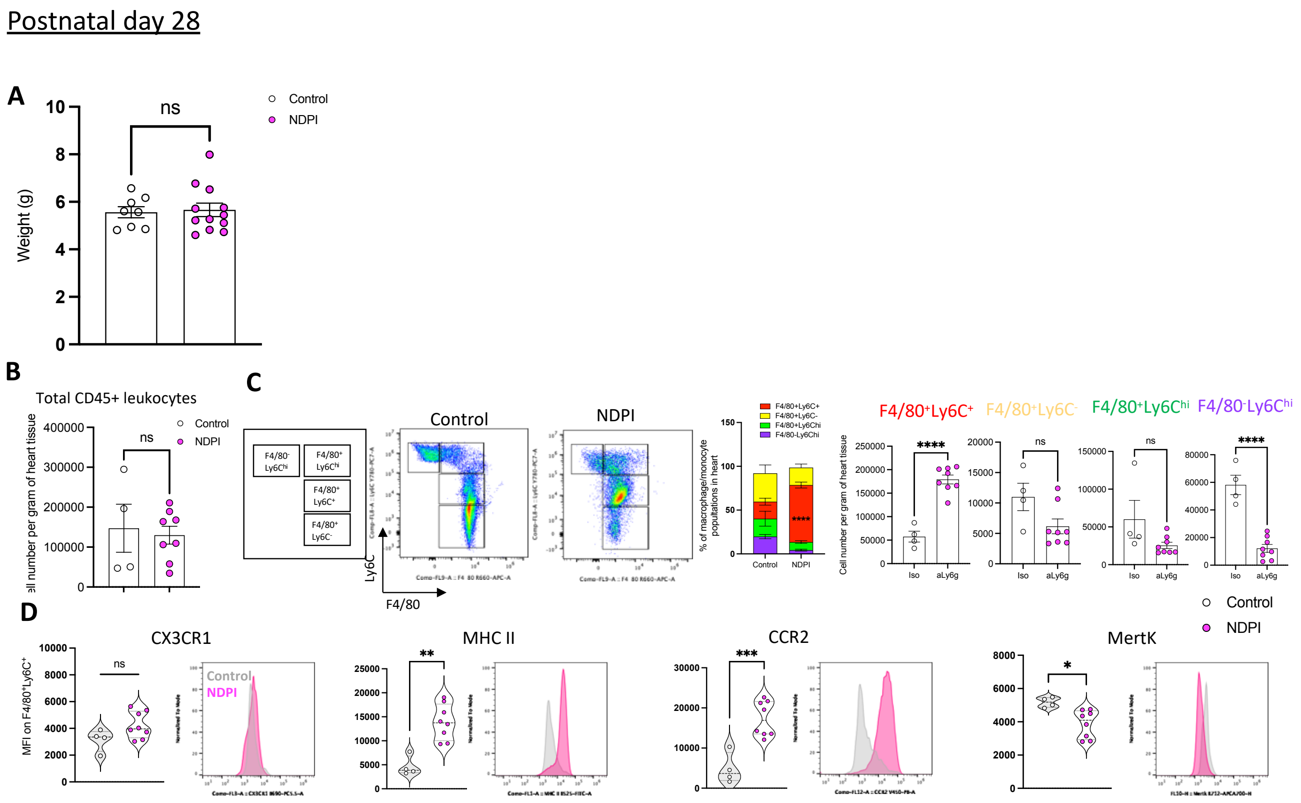


### *Supplementary figure 5*

Neutrophils were depleted at day 4.5 and 7.5 of pregnancy using aLy6G. Offspring of these dams were sacrificed at post-natal day 28 immune composition assessed. Offspring from isotype control treated (white) and neutrophil depleted (NDPI) (pink).

(A) Postnatal day 28 body weight in grams

(B) Graphs showing number of CD45+ leukocytes per gram of heart tissue.

(C) Gating strategy and representative FACS plots of heart cells based on expression of F4/80 and Ly6C. Graphs showing the proportion of various F4/80 Ly6C subpopulations and absolute number per gram of heart tissue.

(D) Graphs showing MFI of CX3CR1, MHCII, CCR2 and MertK expression on F4/80$^{+}$Ly6C$^{+}$ cells from hearts. Representative histogram plots for the indicated marker next to the graphs.

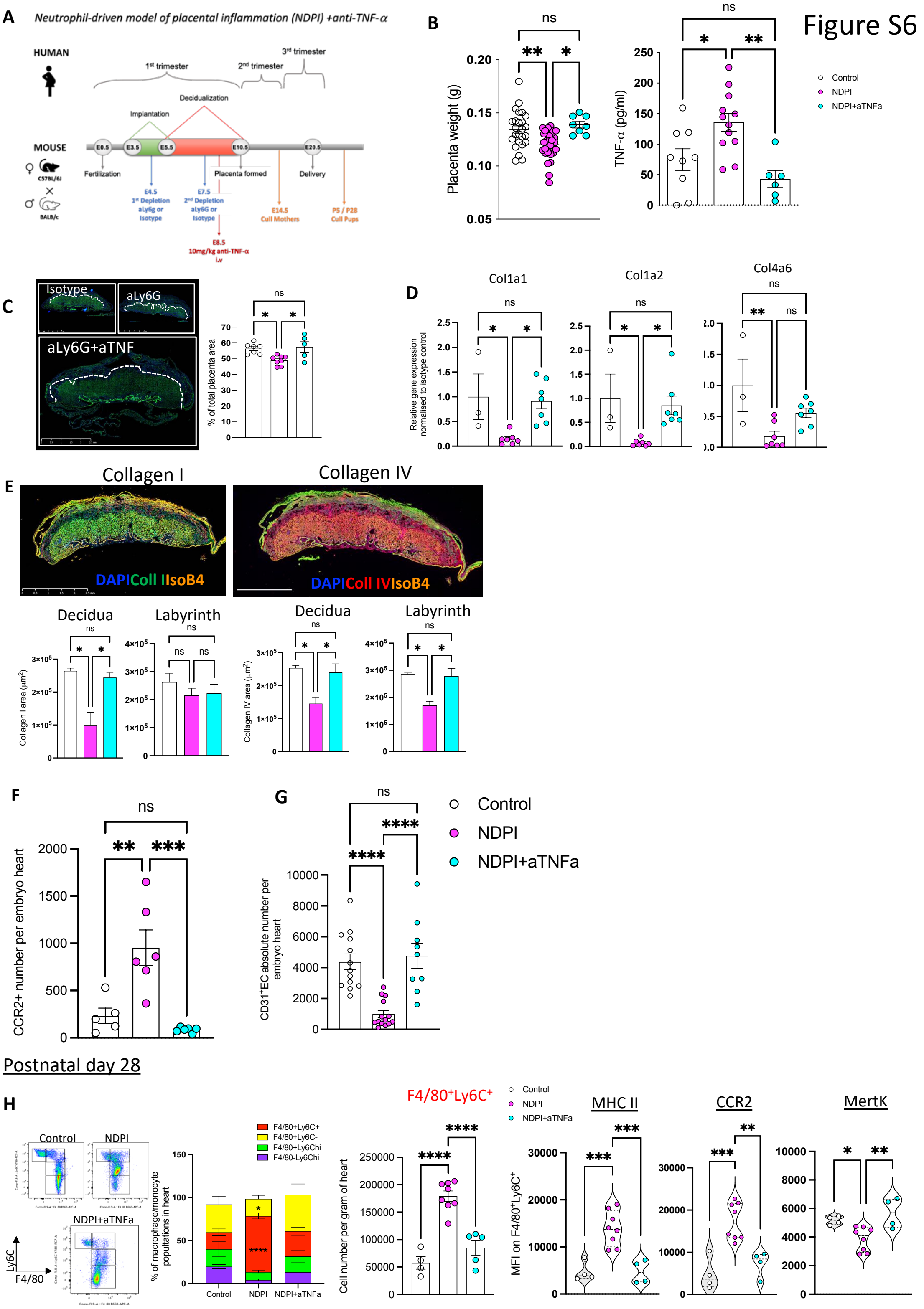

Figure S6
A
Neutrophil-driven model of placental inflammation (NDPI) +anti-TNF-α
HUMAN
1st trimester
2nd trimester
3rd trimester
Decidualization
Implantation
MOUSE
E0.5
E3.5
E5.5
E10.5
E20.5
Fertilization
Placenta formed
Delivery
C57BL/6J
BALB/c
E4.5
1st Depletion
aLy6g or Isotype
E7.5
2nd Depletion
aLy6G or Isotype
E14.5
Cull Mothers
P5 / P28
Cull Pups
E8.5
10mg/kg anti-TNF-α
i.v
B
Placenta weight (g)
TNF-α (pg/ml)
Control
NDPI
NDPI+aTNFa
C
Isotype
aLy6G
aLy6G+aTNF
% of total placenta area
D
Col1a1
Col1a2
Col4a6
Relative gene expression normalised to isotype control
E
Collagen I
Collagen IV
DAPI Coll I IsoB4
DAPI Coll IV IsoB4
Decidua
Labyrinth
Collagen I area (µm²)
Collagen IV area (µm²)
F
CCR2+ number per embryo heart
G
CD31+EC absolute number per embryo heart
Control
NDPI
NDPI+aTNFa
Postnatal day 28
H
Control
NDPI
NDPI+aTNFa
Ly6C
F4/80
F4/80+Ly6C+
F4/80+Ly6C-
F4/80+Ly6Chi
F4/80-Ly6Chi
% of macrophage/monocyte populations in heart
F4/80+Ly6C+
Cell number per gram of heart
MHC II
CCR2
MertK
MFI on F4/80+Ly6C+

***Supplementary figure 6***

(A) Dampening of exaggerated placental inflammation - schematic for experimental design.

(B) Weights of E14.5 placentae. ELISA showing the concentration of TNF-α in placenta digest supernatants. Isotype control treated (white), neutrophil depleted (NDPI) (pink), neutrophil depleted TNFα neutralised (NDPI + aTNFa) (blue).

(C) Immunofluorescent staining of placentae for wheat germ agglutinin (WGA) (green) to show gross structure and cell nuclei with DAPI (blue). White dotted line indicates the separation of the labyrinthine zone/trophoblast area and the outer junctional zone. Graph showing the proportion of the total area of the placentae occupied by the trophoblast zone.

(D) RT-PCR showing the gene expression of Col1a1, col1a2 and col4a6 in placentae from control and NDPI and NDPI + aTNFa pregnancies normalised to control.

(E) Immunofluorescent staining of placentae for Collagen I (green) isolectin b4 (orange) and cell nuclei with DAPI (blue) (left image) or Collagen IV isolectin b4 (orange) and cell nuclei with DAPI (blue) (right image). Graphs showing quantification of the total area of Col I or Col IV expression in the decidua and labyrinth.

(F) Number of $CCR2^{+}$ cells per embryo heart.

(G) Number of $CD31^{+}$ cells per embryo heart.

(H) Experiment as outlined in S1-A. Offspring from treated pregnancies sacrificed at day 28 post-natal. Gating strategy and representative FACS plots of heart cells based on expression of F4/80 and Ly6C. Graphs showing the proportion of various F4/80 Ly6C subpopulations and absolute number per gram of heart tissues. MFI of MHCII, CCR2 and MertK expression on F4/80+Ly6C+ cells from P28 hearts.

Each symbol represents an individual mouse and statistical significance was tested by unpaired Student's t-test. ns = not significant, *p≤0.05, ** p≤0.01, ***p≤0.001 **** p≤0.0001. In all cases, data are mean ± SEM. MFI, median fluorescence intensity
